# Abundant non-inclusion α-synuclein pathology in Lewy body-negative LRRK2-mutant cases

**DOI:** 10.1101/2024.12.20.629583

**Authors:** Nanna Møller Jensen, Zagorka Vitic, Mia R. Antorini, Tobias Bruun Viftrup, Laura Parkkinen, Poul Henning Jensen

## Abstract

Lewy body diseases are common neurodegenerative diseases, including Parkinson’s disease (PD) and dementia with Lewy bodies, which lead to both motor and non-motor symptoms. They are neuropathologically characterized by loss of neuromelanized neurons in the substantia nigra pars compacta and α-synuclein-immunopositive inclusions (Lewy bodies) in several types of neurons in the brain. A fraction of monogenic PD cases, however, represent a conundrum, as they can present with clinical Lewy body disease but do not have Lewy bodies upon neuropathological examination. For LRRK2, the presence or absence of Lewy bodies is not related to any specific mutation in the gene and different clinical presentation and neuropathology can be present even in the same family.

Here, we present the first evidence of widespread α-synuclein accumulation detected with proximity ligation assay (PLA) using the MJFR14-6-4-2 antibody in six Lewy body-negative LRRK2 cases and compare the levels with five patients with neuropathologically-verified Lewy body disease and six healthy controls. We show that non-inclusion aggregated α-synuclein in the form of particulate PLA signal is dominant in the LRRK2 cases, while both Lewy-like and particulate PLA signal is found in late-stage Lewy body disease. Furthermore, LRRK2 cases displayed prominent particulate PLA signal in pontocerebellar tracts and inferior olivary nuclei in the brainstem, which was not seen in idiopathic Lewy body disease cases. These results suggest that Lewy-body negative LRRK2-related PD is not associated with a lack of α-synuclein aggregation in neurons but rather a deficiency in the formation of inclusions.

## Introduction

Lewy body diseases (LBDs), a collective term encompassing Parkinson’s disease (PD), PD with dementia (PDD), and dementia with Lewy bodies (DLB), are common and debilitating neurodegenerative diseases [37, 56]. They feature varying degrees of motor symptoms (resting tremor, rigidity, and bradykinesia) as well as non-motor dysfunctions, such as hyposmia, sleep disturbances, gut dysfunction, depression, and dementia [37, 70]. Neuropathologically, PD is defined by neuronal loss particularly in the substantia nigra pars compacta (SNpc) as well as characteristic proteinaceous inclusions in the neurons of multiple brain regions [18]. The inclusion pathology, encompassing Lewy bodies (LBs) in the soma and Lewy neurites (LNs) in neuronal processes, present in stereotypical patterns used to stage disease severity [4, 12]. Aggregated forms of the neuronal protein α-synuclein, phosphorylated on serine-129, are found in these inclusions in both PD and DLB [18, 49]. Though most PD cases are idiopathic, mutations in a number of genes can cause autosomal dominant and recessive forms of monogenic PD, which constitute approx. 10-15% of cases [8, 17, 37, 58, 59]. Of these, mutations in the leucine rich repeat kinase (LRRK2) have been particularly puzzling, as some LRRK2 patients are clinically diagnosed with PD but do not contain LBs upon autopsy [38, 59, 90]. This has led to discussion whether these LB-negative LRRK2-PD cases also represent a synucleinopathy or a completely different aetiology not associated with α-synuclein aggregation [14, 38, 50, 71, 78].

In this study, we used an α-synuclein aggregate-specific proximity ligation assay (PLA) to study six cases of LB-negative LRRK2 cases. Based on dual antibody recognition followed by a rolling circle signal amplification, α-synuclein PLA has become a predominant strategy for in situ detection of small, oligomeric α-synuclein aggregates, which are undetected by immunohistochemistry [36, 63, 75, 76]. We have previously shown that α-synuclein PLA strongly labels non-inclusion α-synuclein aggregates preceding regular Lewy pathology but has variable detection of LBs and LNs, depending on the antibody and PLA kit vendor (Duolink or Navinci) [36]. In the present study, we used the conformation-specific MJFR14-6-4-2 antibody in the Navinci PLA application that equally detects α-synuclein aggregate pathology whether organized into inclusions (i.e., LBs and LNs) or not (i.e., oligomeric) [36]. This allowed us to assess both the well-characterized Lewy pathology and the less studied non-inclusion (oligomeric) α-synuclein pathology in one tissue section. The PLA staining patterns from LB-negative LRRK2 cases were compared with five idiopathic LBD (Braak stage 6) and six healthy controls. We demonstrate that these LRRK2 cases consistently display α-synuclein PLA-positive non-inclusion pathology and typically at higher levels than regular LBD cases, while most controls only present with very little PLA signal. These results indicate that α-synuclein aggregation is also a characteristic of LB-negative LRRK2 cases. As such, LB-negative LRRK2 cases may be associated with an inability to form LBs and LNs, rather than a lack of α-synuclein aggregate pathology.

## Materials and methods

### Human tissue

Brains from LB-negative LRRK2 cases (n = 6), idiopathic LBD (n = 5), and healthy control (n = 6) were obtained from Parkinson’s UK Brain Bank (PUKBB), Imperial College London, and Oxford Brain Bank (OBB), Nuffield Department of Clinical Neurosciences in University of Oxford, in accordance with approved protocols by the Wales Research Ethics Committee (23/WA/0273) and the Ethics Committee of the University of Oxford (ref 23/SC/0241). All participants had given prior written informed consent for the brain donation. Both brain banks comply with the requirements of the Human Tissue Act 2004 and the Codes of Practice set by the Human Tissue Authority (HTA licence numbers 12275 for PUKBB and 12217 for OBB). Formalin-fixed paraffin embedded (FFPE) tissue sections of 5 µm thickness from the medulla, pons, midbrain, posterior hippocampus, and amygdala were included in the study. Cases included and their clinical, pathological, and genetic data are summarized in Table 1.

**Table 1.** Demographics, clinical and neuropathological characteristics, and LRRK2 mutations in the cohort. AD = Alzheimer’s disease, DLB = dementia with Lewy bodies, LBD = Lewy body disease (PD/PDD/DLB), PD = Parkinson’s disease, PSP = progressive supranuclear palsy. N/A = not available/not assessed. ^a^ Mean ± SD for age at death in the groups: 82.8±12 (LRRK2), 80.7±7.1 (LBD), 82.7±7.7 (control). ^b^ Estimated from neuropathological reports. ^c^ Categorized as low-AD neuropathological change due to CERAD negative stage and Braak tangle stage despite the high Thal phase. ^d^ Typical PSP-tau pathology (tufted astrocytes, coiled bodies, and neuronal tangles). ^e^ Incomplete assessment at autopsy.

| Case ID | Clinical diagnosis | Sex | Age at death (y)<sup>a</sup> | Disease duration (y) | Post-mortem delay (h) | Braak PD stage | LRRK2 mutation(s) | Thal phase (Aβ) | Braak tangle stage (tau) |
| --- | --- | --- | --- | --- | --- | --- | --- | --- | --- |
| **LRRK2 1** | PD | M | 92 | 11 | 16 | LB-negative | M1646T | 1-2<sup>b</sup> | 3 |
| **LRRK2 2** | Healthy control | F | 94 | – | 24 | LB-negative | N2081D | N/A | N/A |
| **LRRK2 3** | Healthy control | F | 87 | – | 76 | LB-negative | M1646T | 5<sup>c</sup> | 2 |
| **LRRK2 4** | PSP | F | 61 | 5 | 43 | LB-negative | N2081D | 0 | 0<sup>d</sup> |
| **LRRK2 5** | PD | M | 84 | 7 | 15 | LB-negative | N2081D, M1646T, P1542S | 0<sup>b</sup> | 1<sup>b,e</sup> |
| **LRRK2 6** | PD | F | 79 | 9 | 32 | LB-negative | G2019S, N2081D | 2 | 1 |
| **LBD 1** | PD | M | 78 | 12 | 32 | 6 | – | 4 | 4 |
| **LBD 2** | DLB+AD | M | 85 | N/A | 96 | 6 | – | 5 | 5 |
| **LBD 3** | PD | M | 80 | 23 | 24 | 6 | – | N/A | 1 |
| **LBD 4** | DLB+AD | F | 79 | N/A | 72 | 6 | – | 5 | 5-6 |
| **LBD 5** | PDD | F | 81 | N/A | 48 | 6 | – | N/A | N/A |
| **Control 1** | Healthy control | F | 88 | – | 100 | 0 | – | 1 | 1 |
| **Control 2** | Healthy control | M | 82 | – | 45 | 0 | – | 2 | 2 |
| **Control 3** | Healthy control | M | 78 | – | 93 | 0 | – | 3 | 2 |
| **Control 4** | Healthy control | M | 70 | – | 43 | 0 | – | 3 | 2 |
| **Control 5** | Healthy control | F | 89 | – | 24 | 0 | – | 1 | 2 |
| **Control 6** | Healthy control | F | 89 | – | 30 | 0 | – | 0 | 2 |

All non-LRRK2 cases had been verified to not contain mutations in LRRK2 or other PD-related genes by genotyping on Illumina’s NeuroX array for neurodegenerative diseases, encompassing more than 24,000 neurodegeneration-specific variants in addition to the standard Illumina exome content of approx. 240,000 variants [52, 53].

### Proximity ligation assay

Conformation-specific α-synuclein antibody MJFR14-6-4-2 (MJF-14; Abcam, #ab214033, 1 mg/mL) was conjugated to complementary NaveniLink proximity probes (Navinci, #NL.050) according to manufacturer’s instructions. Briefly, 10 µL of Modifier was added to 100 µL antibody, before mixing with the lyophilized oligonucleotides (Navenibody 1 or Navenibody 2) and incubation overnight at room temperature (RT). 10 µL of Quencher N was then added and incubated for 15 minutes at RT, where-after conjugated antibodies were stored at 4°C.

The PLA staining was conducted using NaveniBright HRP kits using DAB as the chromogen (Navinci, #NB.MR.HRP.100) and counterstained with a haematoxylin-based nuclear dye. FFPE sections were deparaffinized and rehydrated in decreasing alcohol series, before antigen retrieval of microwaving for 2x 5 min in sodium citrate (pH 6, DAKO, #S1699). Endogenous peroxidase activity was quenched in 0.3% hydrogen peroxide in PBS, after which samples were blocked in Navinci blocking buffer with supplement 1 for 1 hour at 37°C. PLA-conjugated MJF-14 was diluted 1:10,000 in antibody diluent with supplement 2 and incubated with the samples overnight at 4°C. After washing off unbound antibody, samples were incubated in freshly prepared Reaction 1 (1x Buffer 1 diluted in nuclease-free water and supplemented with Enzyme 1) for 30 min at 37°C. Subsequently, Reaction 2 was similarly prepared (1x Buffer 2 diluted in nuclease-free water and supplemented with Enzyme 2) and applied for 1 hour at 37°C. Sections were then incubated in HRP detection solution for 30 min at RT, before signal development for 5 min at RT. Finally, sections were briefly counterstained in Navinci Nuclear stain and staining developed in tap water, before a quick dehydration in 100% isopropanol and mounting with VectaMount Express Mounting Medium (VectorLabs, #H-5700).

### Immunohistochemistry

Immunohistochemistry was performed as previously described for the Syn-O4 antibody [1, 83]. In brief, FFPE sections were deparaffinized and rehydrated as for PLA, followed by antigen retrieval for 15 min in 80% formic acid at RT. Endogenous peroxidase activity was quenched in 3% hydrogen peroxide in PBS, after which samples were blocked in 10% foetal bovine serum. Mouse monoclonal antibody Syn-O4 against aggregated α-synuclein [83] was diluted 1:5,000 and incubated on the sections overnight at 4°C. Sections were incubated with HRP-conjugated anti-mouse secondary antibody as part of the REAL EnVision detection system (DAKO, #K5007) for 1 hour at RT. Signal was visualized with 3,3ʹ-diaminobenzidine (DAB) and sections were counterstained with haematoxylin. Finally, sections were dehydrated in increasing alcohol concentrations, cleared in xylene, and mounted with DPX mounting medium (Sigma, #06522).

### Imaging and image analysis

Tissue sections were imaged at X20 magnification on an Olympus VS120 slide scanner and stored in the vsi file format. Digitalized images were opened in QuPath [3], stain vectors estimated using the auto function, and the entire tissue regions were outlined by thresholding on averaged channel values. Specific regions of interest were annotated according to The Brain Atlas and specific articles pertaining individual subregions as indicated in Suppl. Fig. 3 [2, 29, 72, 86, 89].

5 total slide scans were selected to optimize parameters for detection of signals for quantitative image analysis. These cases (LBD 1 medulla, LBD 1 pons, LRRK2 6 midbrain, LRRK2 1 posterior hippocampus, and LRRK2 4 amygdala) encompassed different composition and densities of the various signals found across the cohort. Signals were divided into particulate PLA signal (not associated with inclusions) and Lewy-like PLA signal (signal found in structures resembling LBs and LNs) and quantified as area coverage (%) to allow a direct comparison between the two measures. Any neuromelanin in the various regions of interest was manually outlined and excluded from the analysis to avoid the algorithm mistaking it for PLA signal. Then, PLA signals were detected in a three-step protocol. First, all PLA signal was defined using a simple thresholder on the DAB channel with Gaussian pre-filtering with a smoothing sigma of 0.5 and a threshold of 0.1. Next, Lewy-like PLA signal was determined using a simple thresholder from the average channel intensity with Gaussian pre-filtering of 1.0 and a threshold of 80, where anything below the threshold corresponded to Lewy-like PLA. Finally, particulate PLA was defined as signal positive on the DAB thresholder but negative on Lewy pathology thresholder. The image analysis strategy is illustrated in Suppl. Fig. 1.

### Statistical analysis

To compare levels of particulate and Lewy-like PLA between groups, we ran univariate analyses adjusting for age and sex using SPSS v.29.0.1.0 (IBM). Other potential covariates (post-mortem delay, Braak tau stage, and Aβ Thal phase) were assessed for their influence on the model and generally not included. Data were tested for assumptions of normal distribution of residuals using Shapiro-Wilk’s test, linear relation of covariates to the dependent variable, homogeneity of regression slopes, homoscedasticity, and homogeneity of variances using Levene’s test. In cases where assumptions of normality of residuals and/or homogeneity of variances were violated, the dependent variable was log-transformed, and data were re-tested to assure that they now fulfilled the assumptions. Correction for multiple comparisons in the analyses was done using Bonferroni’s post-test. P-values under 0.05 were considered significant (* p<0.05, ** p<0.01, *** p<0.001), though values close to significance were also indicated, as the small group sizes limit the chance of reaching statistical significance. All graphs were compiled using GraphPad Prism 10, and data were plotted as mean ± SEM unless otherwise mentioned.

## Results

The study included six LRRK2 cases, all of which were LB-negative on standard neuropathological assessments. Three of the cases had a clinical diagnosis of PD, one a diagnosis of progressive supranuclear palsy (PSP, neuropathologically confirmed), and two cases had no clinical diagnoses of neurodegenerative disease. These were compared with five idiopathic LBD cases, all of which were Braak stage 6, and six healthy controls, as based on standard clinicopathological assessments. None of the idiopathic LBD cases or the controls harboured any mutations in LRRK2 or other LBD-related genes.

Clinical, demographic, genetic, and neuropathological information on the cases is summarized in Table 1.

From each case, FFPE sections from five brain regions were stained with an α-synuclein aggregate PLA using the MJF-14 antibody (Fig. 1a). The PLA staining produced two distinct patterns of signal, 1) punctate PLA particles that appeared to locate to the neuropil and were not associated with neuronal inclusions, called “particulate PLA” (Fig. 1b, top) and 2) a Lewy-like PLA signal strongly resembling the LBs and LNs stained by regular IHC (Fig. 1b, bottom). We have previously confirmed by immunofluorescent double-staining with Navinci MJF-14 PLA and pSer129-α-synuclein that the Lewy-like PLA-staining indeed co-localizes with the pSer129-α-synuclein labelled LBs [36]. The two patterns of signal were distinguished based on their colour and intensity, as the Lewy-like PLA signal (formed by coalescing of highly abundant PLA particles) appeared intensely reddish black (Suppl. Fig. 1). While Lewy-like PLA signal is primarily found in the neuronal somas (LBs) and processes (LNs), the particulate PLA signal is often located outside the neuronal somas in the neuropil (Fig. 1b). Although the exact subcellular distribution of PLA signal is not fully studied, it has been demonstrated in presynaptic terminals, neuronal somas, glial cells, and neuronal processes [36, 75, 76].

**Fig. 1.**
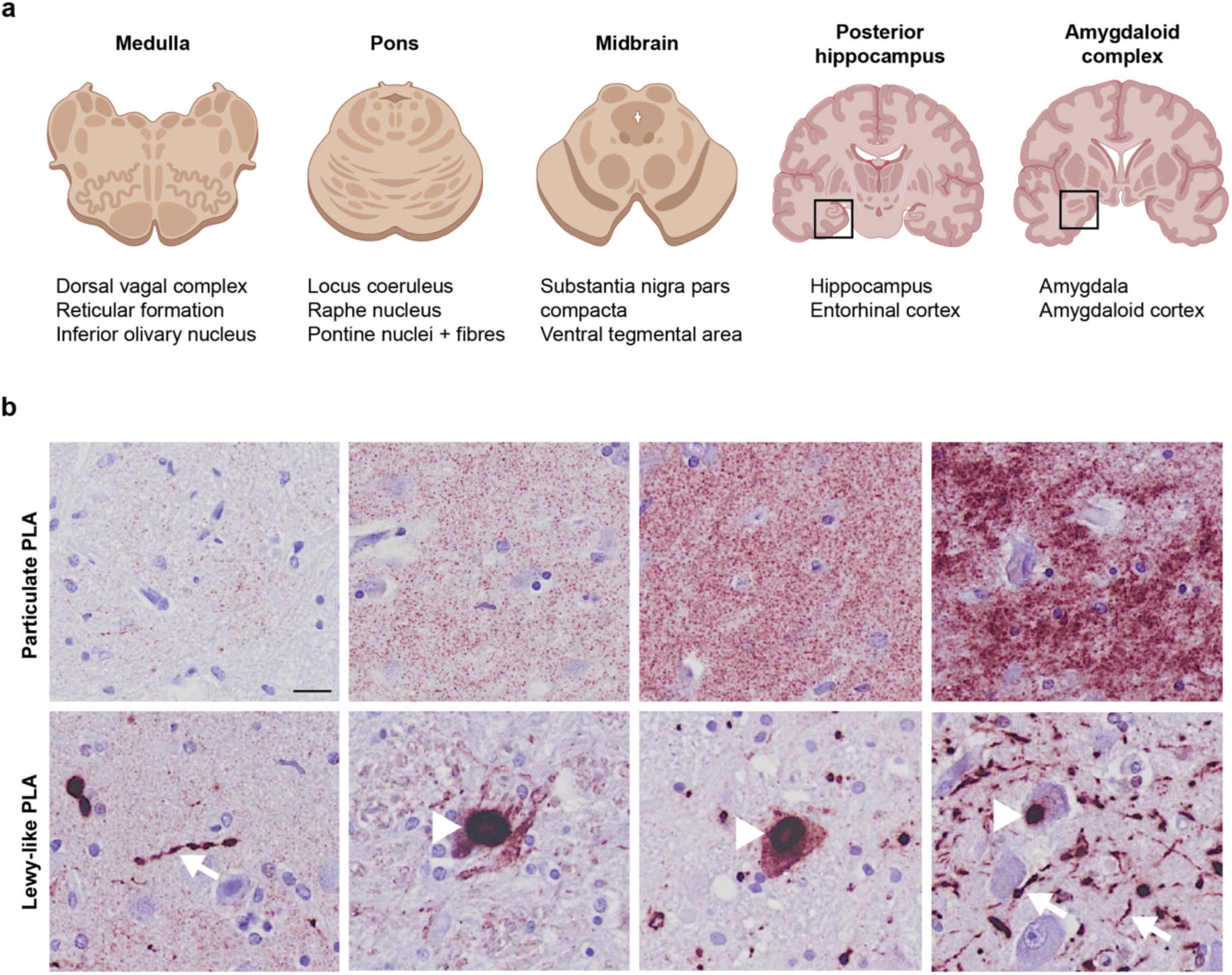
Regions of interest and PLA terminology. **a**) Five brain regions (medulla, pons, midbrain, posterior hippocampus, and amygdala) were included for each case to allow comparative quantitative analyses of PLA signals. See Suppl. Fig. 3 for delineation of the listed subregions in three of the cases included in this study. **b**) The PLA staining yielded two distinct patterns of signal: the particulate PLA not associated with inclusions (top panels) with increasing signal density from left to right and a Lewy-like PLA staining (bottom panel) with strong morphological similarity to LBs and LNs as stained by standard IHC. Presumed LBs are indicated by arrowheads, while examples of putative LNs are highlighted with arrows. Scale bar = 20 µm (applies to all images). Particulate and Lewy-like PLA were analysed separately in all sections and reported as area covered (%).

### LRRK2 cases show abundant particulate PLA, while both particulate and Lewy-like PLA is found in LBD cases

As expected from the case inclusion criteria, Lewy-like PLA signal was almost exclusively present in the Braak stage 6 LBD cases, where it was found in all regions analysed (Fig. 2a, c). The Lewy-like PLA encompassed both LBs and LNs, the latter of which were abundant in e.g. the CA2 of the hippocampus (Fig. 2a). Virtually no Lewy-like PLA signal was detected by automated quantification (see Suppl. Fig. 1) in either LB-negative LRRK2 cases or healthy controls (Fig. 2c). In contrast, particulate PLA signal was predominant in the LRRK2 cases, of which all six contained the signal (Fig. 2a-b), despite a general lack of distinctive staining by Syn-O4-immunohistochemistry (IHC) against aggregated α-synuclein (Suppl. Fig. 2b). Occasional punctate axonal varicosities were detected by Syn-O4 IHC in three of the LRRK2 cases, though these were much less prominent than the particulate PLA (Suppl. Fig. 2c). Especially the posterior hippocampus and amygdala sections of the LRRK2 cases contained high levels of particulate PLA, with an average area coverage of 8-10% (Fig. 2b).

**Fig. 2.**
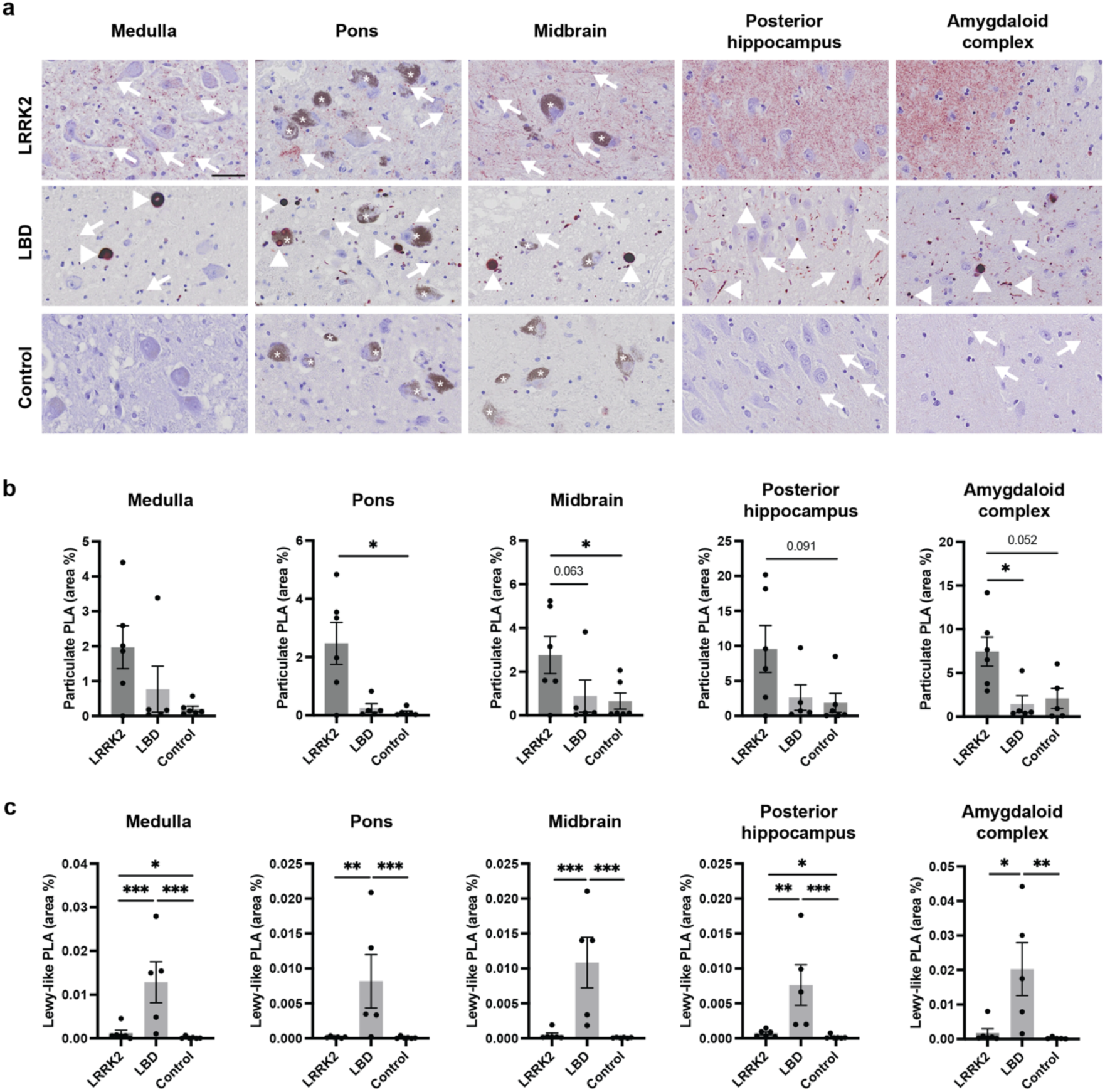
Particulate PLA, Lewy-like PLA, and LB densities in LRRK2 LB-negative cases, regular LBD, and non-neurodegenerative controls. **a**) Overview of PLA staining appearance in medulla (around DMV), pons (around LC), midbrain (around SNpc), posterior hippocampus (around CA2), and amygdaloid complex (in the amygdala). Scale bar = 50 µm (applies to all images). **b-c**) Quantitative analysis performed in the entire tissue sections showing particulate PLA area coverage (%, **b**) and Lewy-like PLA area coverage (%, **c**) for the three groups. All graphs show mean ± SEM with data points for individual cases. Univariate analyses adjusting for age and sex followed by Bonferroni’s multiple comparison test. * p<0.05, ** p<0.01, *** p<0.001. CA2 = cornu ammonis 2 of the hippocampus, DMV = dorsal motor nucleus of the vagus, LC = locus coeruleus, SNpc (substantia nigra pars compacta).

In addition to its presence in the LRRK2 cases, some particulate PLA signal was also found in LBD cases and two of controls (Fig. 2a-b). In general, the LB-negative LRRK2 cases contained 2.5-10 times more particulate PLA signal than the LBD cases when analysing the entire tissue sections, though group differences only reached statistical significance for a couple of regions, when correcting for multiple comparisons with Bonferroni (Fig. 2b). This indicates that non-inclusion α-synuclein aggregates are more abundant in the LB-negative LRRK2 cases than LBD cases or controls, though larger groups are needed to determine more precise effect sizes and achieve statistical significance in individual brain regions.

Furthermore, even in the LBD cases, the particulate PLA signal was much more abundant than the Lewy-like PLA signal, in line with previously published studies using α-synuclein PLA in LBD cases [36, 75, 76]. This was also apparent when comparing the MJF-14 PLA with Syn-O4 IHC, which showed that especially particulate PLA signal outside the cell bodies, but also non-inclusion accumulations in the neuronal somas, were preferentially detected with PLA but not with IHC (Suppl. Fig. 2a). Of note, the ratio between particulate and Lewy-like PLA increased with more rostral sections, with an average area coverage from 41-fold higher in medulla to more than 100-fold higher in posterior hippocampus and amygdala sections (Fig. 2b-c, Suppl. Table 1). Based on the limited cohort in this study, we could not conclude whether this difference in signal ratio was related to region-specific propensities for forming LBs (brainstem versus limbic/cortical), a rostral progression of disease, or other yet unknown causes.

### LRRK2 cases display varying amounts of particulate PLA signal in nuclei and regions typically affected in PD but also in unusual regions

To investigate the distribution of particulate and Lewy-like PLA in more detail, we outlined a number of subregions in each tissue section for quantitative analysis (see Suppl. Fig. 3 for representative overviews and delineation of regions). In general, the LRRK2 cases showed particulate PLA signal in most classical PD regions such as the dorsal motor nucleus of the vagus (DMV), locus coeruleus (LC), SNpc, and ventral tegmental area (VTA) (Fig. 3). The idiopathic LBD cases typically showed similar levels of particulate PLA in these nuclei and regions, in addition to the prominent LBs (Fig. 3, see quantifications in Suppl. Fig. 4).

**Fig. 3.**
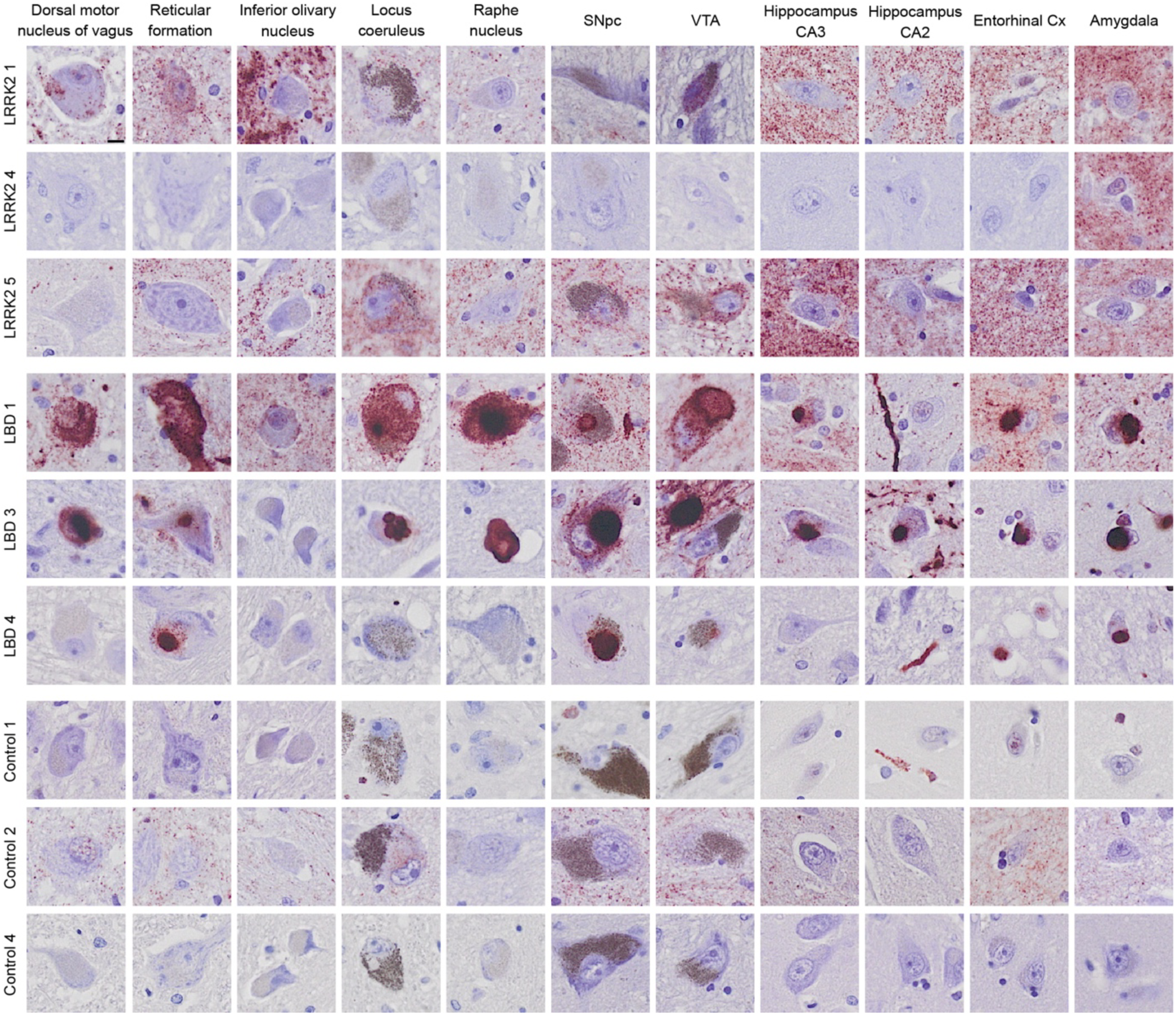
PLA signal in specific brain regions from selected cases. Specific brain regions were identified from the medulla (dorsal vagal complex, reticular formation, and inferior olivary nucleus), pons (locus coeruleus and raphe nucleus), midbrain (substantia nigra pars compacta, SNpc, and ventral tegmental area, VTA), posterior hippocampus (CA3, CA2, and entorhinal cortex) and amygdala. Three cases were displayed from each group, encompassing most of the variation in signal densities in each group. Scale bar = 10 µm (applies to all images). Note that LRRK2 4 was almost entirely depigmented and had substantial neuronal loss in several regions, incl. SNpc. See also semi-quantitative overview particulate versus Lewy-like PLA in all cases in Fig. 6.

However, the particulate PLA in LRRK2 cases was especially prominent in regions not typically affected in PD, such as the inferior olivary nucleus (ION) (Fig. 4) and the basilar pons (Fig. 5). These regions were not stained by IHC for aggregated α-synuclein using the Syn-O4 antibody, except for the occasional weak brown background, which did not coincide with PLA staining (Suppl. Fig. 5). In five of the six LRRK2 cases, the ION contained noteworthy particulate PLA signal, while only one of five LBD cases displayed signal in this region (Fig. 4). Of note, where the particulate PLA in the LRRK2 cases was confined to the neuropil, sparing the cell bodies of ION neurons, LBD 1 showed particulate PLA signal in both the neuropil and the ION neurons (Fig. 4b, arrows). In the basilar pons, the transverse, ponto-cerebellar fibres were highly positive for particulate PLA in the same five of six LRRK2 cases, along with varying positivity in the pontine nuclei (Fig. 5a-b). The longitudinal fibres, which encompass the ascending and descending corticospinal and corticonuclear fibres, only contained negligible levels of particulate PLA signal (Fig. 5a-b). In the LBD cases, in comparison, the pontine nuclei and transverse, pontocerebellar fibres only contained low levels of signal, whether particulate or Lewy-like PLA (Fig. 5a, c). As for the LRRK2 cases, the longitudinal fibres were generally negative for any PLA signal in the LBD cases as well, except for LBD 5 in which some Lewy-like PLA signal was found (Fig. 5c, arrows). Quantification of the particulate PLA in the transverse pontocerebellar fibres indeed showed that the LRRK2 cases contained significantly more signal than either LBD cases or controls (Suppl. Fig. 4).

**Fig. 4.**
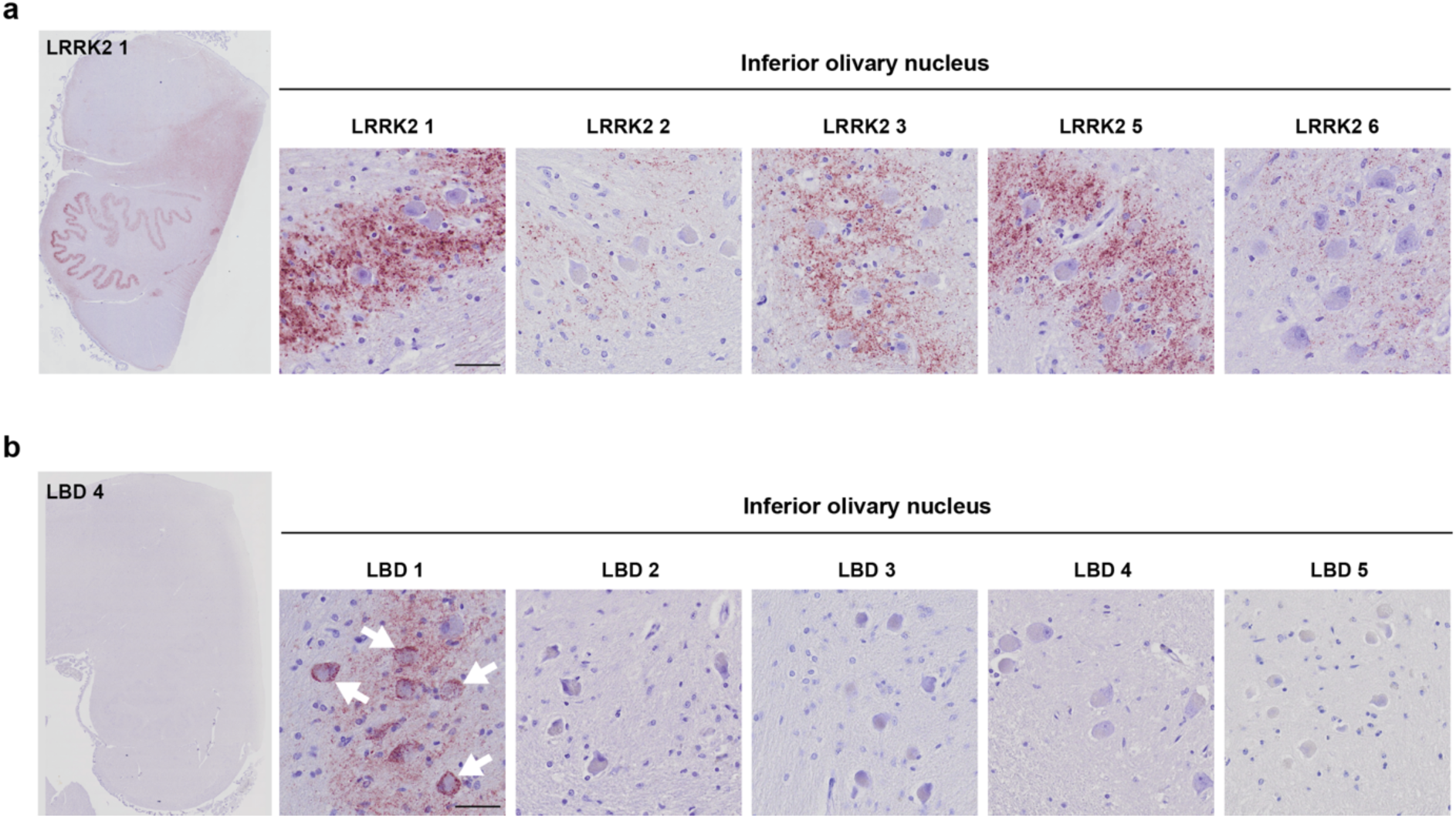
Prominent PLA signal in the inferior olivary nucleus of LRRK2 cases. **a**) LRRK2 cases displayed particulate PLA signal in the inferior olivary nucleus, with signal predominantly present in the neuropil surrounding the neuronal cell bodies. Note that LRRK2 4, which contained no PLA signal in the medulla, was left out of the figure. **b**) Only LBD 1 of the LBD cases displayed PLA signal in the inferior olivary nucleus. Note that in this case, signal is not confined to the neuropil but also found in the neuronal cell bodies. Scale bars = 50 µm. See also semi-quantitative overview particulate versus Lewy-like PLA in all cases in Fig. 6.

**Fig. 5.**
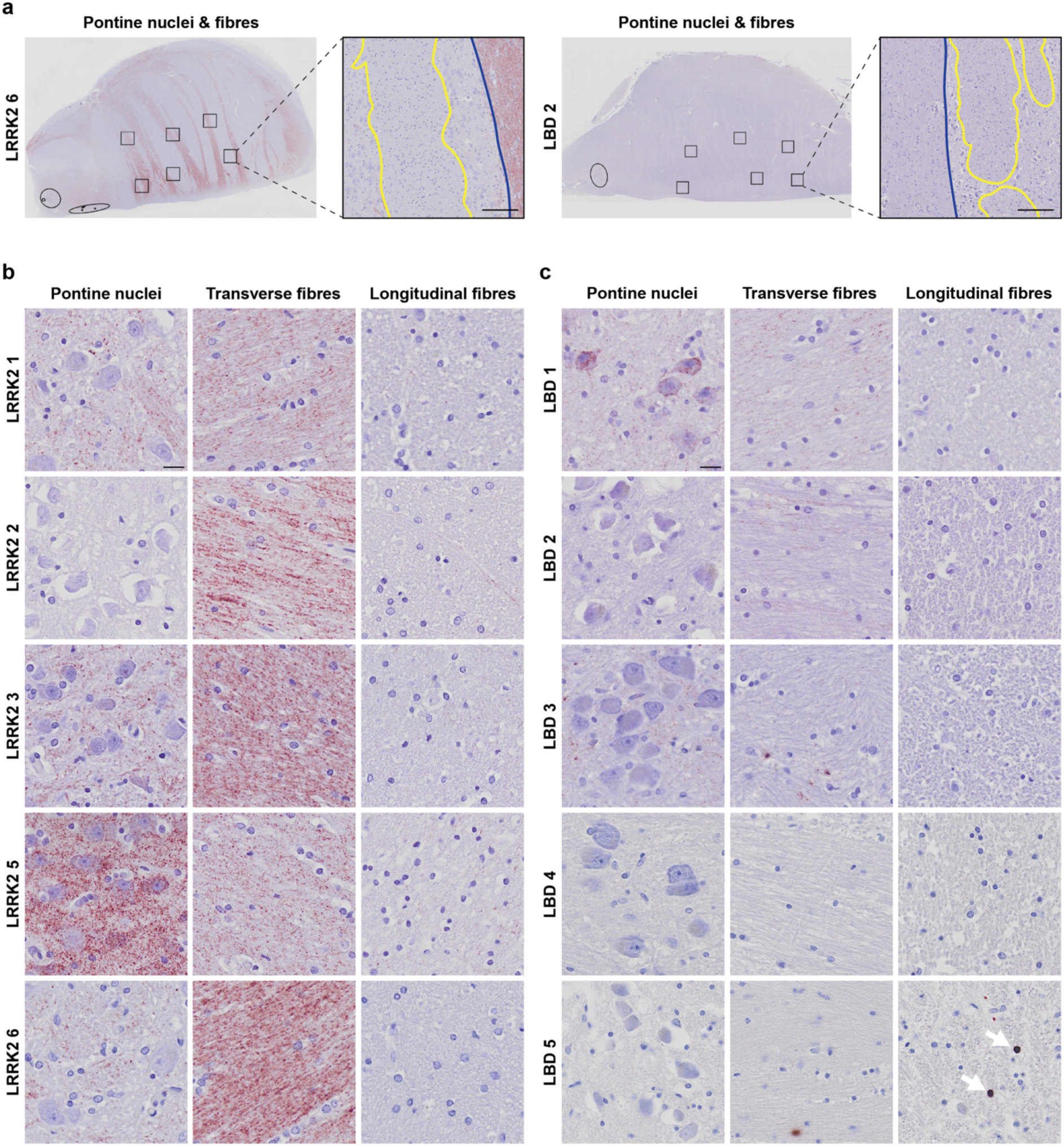
PLA in pontine nuclei, transverse and longitudinal fibres of the basilar pons. **a**) Particulate and Lewy-like PLA signal was assessed in six ROIs in the basilar pons of all cases as indicated in the overviews from one LRRK2 case (left) and one LBD case (right). Magnified panels show transverse fibres (outlined in blue) and longitudinal fibres (outlined in yellow), while in-between areas correspond to pontine nuclei. Scale bars = 200 µm. **b**) LRRK2 cases displayed dense particulate PLA signal in the transverse (pontocerebellar) fibres and to some degree in the pontine nuclei. Note that LRRK2 4, which was negative in the pons, was left out of the figure. Scale bar = 20 µm (applies to all images). **c**) LBD cases displayed much less signal in pontine nuclei and transverse fibres than the LRRK2 cases. One case (LBD 5) presented with some Lewy-like PLA signal in the longitudinal fibre bundles (arrows). Scale bar = 20 µm (applies to all images). See also semi-quantitative overview particulate versus Lewy-like PLA in all cases in Fig. 6 and quantifications in Suppl. Fig. 4.

In addition to the prominent brainstem pathology, five out of six LRRK2 cases displayed high levels of particulate PLA in hippocampus and entorhinal cortex, and all cases in amygdala-associated cortex (Fig. 3, Suppl. Fig. 3-4). In the hippocampus, particulate PLA signal was found in all hippocampal sub-regions (dentate gyrus as well as CA1-3), though the specific distribution varied between cases. In entorhinal and amygdala-associated cortex, particulate PLA signal was present in both the superficial (layers 1-3) and deep cortical layers (layer 5-6), without any obvious difference in signal density. This was observed for both the LRRK2 cases and the LBD cases, though LBs were selectively accumulated in the deep cortical layers, in line with previous reports from us and others [11, 30, 36].

Healthy control cases generally displayed low particulate PLA signal, though two cases showed some abundance around specific neuronal populations in the DMV, SNpc, and amygdala (see Controls 2 and 6, Fig. 3 and semi-quantitative overview in Fig. 6).

**Fig. 6.**
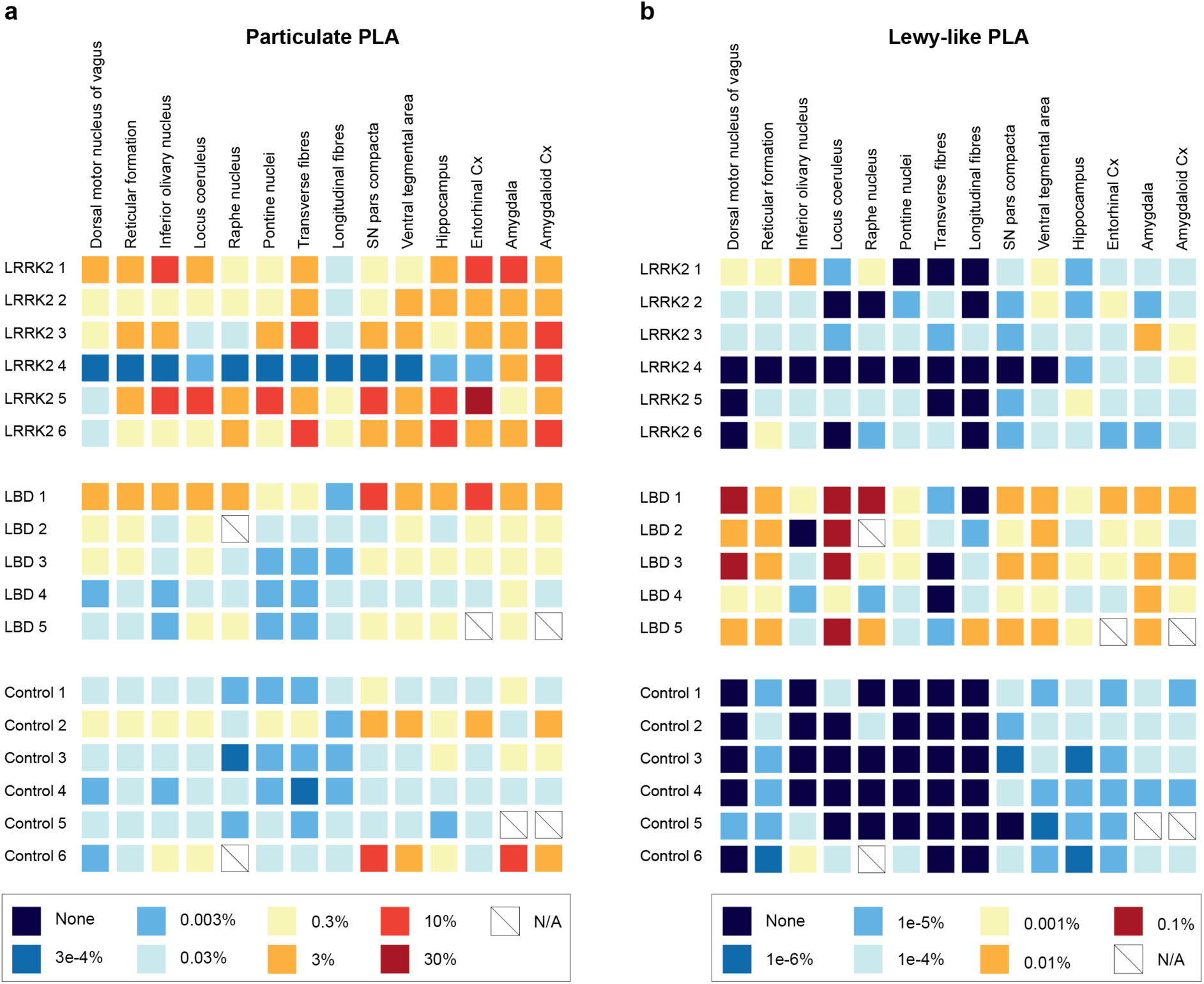
Graphical summary of particulate and Lewy-like PLA across all cases and nuclei/regions examined. Based on the quantifications of area coverage, semi-quantitative scales were made for particulate PLA (**a**) and Lewy-like PLA (**b**). Note that the same colour in **a** and **b** denotes a 300-fold difference in area %. LBD cases stand out with a high density of Lewy-like PLA across the regions examined, while LRRK2 cases display the highest particulate PLA density. N/A, not available.

As already seen in the quantifications in Fig. 2 and Suppl. Fig. 4, there was quite a lot of variation in signal density (both particulate PLA and Lewy-like PLA) within the LRRK2 and LBD case groups, respectively. No single case or mutation (for the LRRK2 cases) was consistently associated the highest signal (particulate or Lewy-like PLA) across the analysed regions for each group. One of the LRRK2 cases (LRRK2 4) stood out amongst the cases with profound particulate PLA in the amygdala alone, while all other regions were unaffected (Fig. 3, Fig. 6a). All of the other LRRK2 cases had significant particulate PLA in all brain sections examined, although the densities and regional distributions differed. Curiously, the LRRK2-mutation present in LRRK2 4 (N2081D) was also present in LRRK2 2 (and in LRRK2 5 and 6 along with additional mutations), indicating that the unique staining pattern in LRRK2 4 cannot be solely ascribed to the N2081D LRRK2-mutation (Table 1). Alternatively, either the clinical diagnosis of progressive supranuclear palsy (PSP), which was the only tauopathy diagnosis in the cohort, the relatively young age at death compared to the other LRRK2 cases, or other factors may affect the presence of particulate PLA.

### Neuronal loss may explain some of the variation in PLA signal density between cases

One of the factors likely affecting the signal densities observed in the LRRK2 and LBD cases is the neuronal loss in multiple brain regions, which is well-characterized for PD [18, 21, 23, 51]. In a few cases, in particular LBD 4, brainstem nuclei such as the LC and SNpc were almost entirely depigmented and only few neurons were left. Though we did not specifically assess the cell loss, significant loss of cells in these nuclei may have resulted in a lower signal density, both for the particulate and the Lewy-like PLA. Indeed, LBD 4 for example, presented with lower signal density in the LC than the other LBD cases and also contained substantially fewer neurons in this region upon closer examination (Fig. 6 and Fig. 7). As a corroborating feature, supporting neuronal cell loss, we also observed extra-neuronal neuromelanin in several of the cases [18] (Fig. 7, arrowheads). Further studies to determine the exact relationship between PLA signal, neuronal cell loss, and/or neuromelanin would require larger cohorts, though it would seem logical that substantial neuronal cell loss would result in less neuronal pathology.

**Fig. 7.**
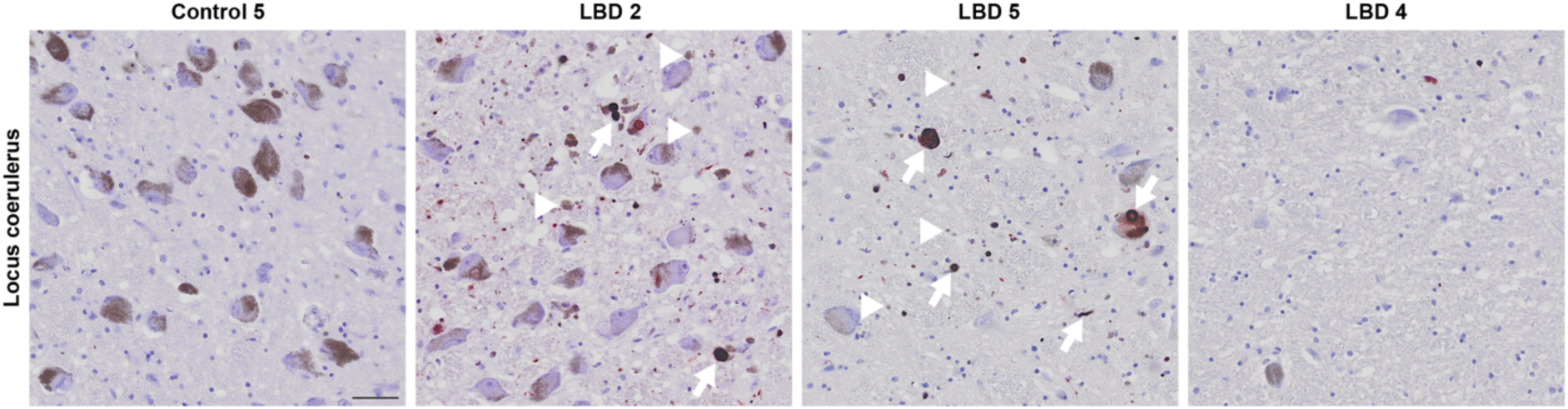
Cell loss and extra-neuronal neuromelanin in the locus coeruleus of LBD cases. Locus coeruleus displayed varying degrees of loss of its noradrenergic, neuromelanin-containing neurons in the LBD cases, as illustrated with a progressive fall in the density of melanized neurons (left to right). Extra-neuronal neuromelanin was apparent in some cases (arrowheads in LBD 2 and 5), evincing neuronal loss. PLA signal density was also lower in LBD cases with extensive neuronal loss (arrows indicate examples of Lewy-like PLA). Scale bar = 50 µm.

## Discussion

In recent years, several studies have demonstrated previously undetected α-synuclein aggregate pathology not organised into LBs in PD, DLB, and multiple system atrophy (MSA) using α-synuclein PLA [36, 63, 75, 76]. With their signal amplification creating a high signal-to-noise ratio, the PLAs are capable of detecting aggregate species with low epitope density, which enables detection of, e.g., α-synuclein oligomers. Oligomers and other small α-synuclein aggregates are generally not detected by IHC, as their epitope density is so low that – even if the antibodies label them – they are indistinguishable from physiological α-synuclein species or deemed as background [5]. Some studies have used proteinase K pretreatment to degrade physiological α-synuclein before IHC, demonstrating abundant synaptic α-synuclein aggregates that appear comparable to PLA-detected non-inclusion aggregates [41, 54, 73, 80]. Nevertheless, the proteinase K pretreatment also affects tissue integrity and is likely to degrade α-synuclein oligomers with a low beta-sheet content [5]. How the α-synuclein PLA signal corresponds to these proteinase K resistant synaptic aggregates and whether oligomers with low beta-sheet content contribute to the α-synuclein PLA signal is currently unknown.

In the present study, we used the Navinci version of the recently published MJF-14 PLA [36], which labels both inclusions of Lewy-type morphology as well as the non-inclusion aggregates. Though the exact nature of the α-synuclein species detected by the PLA is not yet determined, the use of the aggregate-specific MJF-14 antibody should ensure that only pathological α-synuclein species are detected [7, 36, 66]. With this assay, we present the first evidence of a broad accumulation of α-synuclein aggregates not organised into inclusions in multiple brain regions of LRRK2 cases without LBs. Of the six cases examined, five of them presented with significant particulate PLA signal in all the five brain regions examined, but the exact distribution and density differed between the various cases and brain regions investigated. Though most of the LRRK2 mutations in our cases (Table 1) are not traditionally listed as pathogenic/PD-related, there is evidence suggesting a link between PD and mutations in both M1646T and N2081D [13, 42, 64, 79]. Some studies have also shown increased LRRK2 kinase activity with M1646T and N2081D mutations compared to wildtype LRRK2, though to a lower degree than with G2019S LRRK2 [35, 39, 62]. Lastly, M1646T and N2081D mutations in LRRK2 were quite prevalent among PD-diagnosed LRRK2 cases with Lewy pathology in the Parkinson’s UK brain bank – more so than G2019S or R1441C/G/H mutations. With the low penetrance of LRRK2 mutations for development of PD (estimated 25-40% for G2019S LRRK2 [13]), it is clear that there is still much to uncover regarding the pathogenicity of LRRK2 mutations and their interaction with other modulators of disease penetrance.

Curiously, several of the LRRK2 cases did not show particularly dense PLA staining in the brainstem regions such as DMV, LC, and SNpc. Instead, we detected a noteworthy affection of inferior olivary and pontine nuclei as well as pontocerebellar fibres, raising the speculation whether this is a general feature of LRRK2 cases. These regions are generally implicated in MSA rather than the LBDs, though sparse Lewy pathology has been reported in some PD and DLB cases [74, 85]. If this observation is shown to be a common characteristic in LRRK2-mutant cases, it may suggest a slight cerebellar dysfunction in these patients, which should then be investigated clinically. In general, the LRRK2 cases presented with a higher degree of particulate PLA signal than both the regular LBD cases and the non-neurodegenerative controls, though one LBD case did show particulate PLA signal at levels similar to most of the LRRK2 cases.

The striking prevalence of particulate PLA in the LRRK2 cases suggests that their lack of LBs does not reflect an absence of disease-associated α-synuclein aggregates but rather an inability to form the characteristic aggregate-containing inclusions. Though knowledge of the mechanisms involved in LB formation is limited, LBs are clearly complex structures, and their formation appears to be a specific cellular response, which might indeed be perturbed in some cases [20, 43, 46, 55, 77, 81, 84]. LRRK2 has mechanistically been associated with vesicular trafficking and autophagy, and mutation-associated perturbations of proteostasis could thus represent a link to the absence of LBs [28, 45, 69]. However, since the absence of LBs is not linked to any specific LRRK2-mutation and varies even within the same family, LRRK2 is not the sole player in the perturbation of LB formation [38, 59, 90]. Consequently, further studies to determine whether such non-inclusion α-synuclein aggregate pathology is a general feature of PD patients with mutations in LRRK2 or in Parkin (PRKN), who also frequently present without LBs, are strongly motivated [19].

Curiously, a recent study using α-synuclein seed amplification assay (SAA) on cerebrospinal fluid (CSF) from more than 1100 participants in the Parkinson’s Progression Marker Initiative (PPMI) cohort showed that SAA-positivity was significantly lower among LRRK2 PD than idiopathic PD patients [78]. At the time of the study, neuropathological examination was only available for a small subset of cases, including one SAA-negative LRRK2 carrier, who did not contain Lewy pathology. In contrast, all cases with a positive SAA showed typical Lewy pathology upon neuropathological examination [78]. At present, the correlation between SAA-positivity and presence of α-synuclein PLA signal has not been studied, though such studies would be highly valuable. A recent study, however, did examine the relation between CSF SAA-positivity and regional presence of Lewy pathology, demonstrating significantly lower CSF SAA-positivity in cases with early (Braak stages 1-3) or focal pathology (e.g., amygdala-predominant) [6]. Moreover, they found SAA-positivity in brain homogenates from multiple brain regions without accompanying Lewy pathology, indicating a seeding activity of smaller or less organized α-synuclein aggregates [6]. As such, the lack of CSF SAA-positivity in some of the LRRK2 cases in the PPMI cohort does not necessarily reflect an absence of α-synuclein aggregates but could also be caused by predominantly focal pathology, which is not reflected in the cerebrospinal fluid [6].

Alternatively, pathology consisting of non-inclusion (presumably oligomeric) α-synuclein may have a lower seeding efficiency than mature α-synuclein fibrils in the SAA, in line with evidence from experimental models [47, 57]. Additional SAA analyses on homogenates from various brain regions could thus be an important next step to assess the presence of seeding-competent α-synuclein aggregates, even in the absence of Lewy pathology. Combined with α-synuclein PLA staining in post-mortem cases from the upcoming follow-up studies on the PPMI cohort, this will enhance our understanding of the LRRK2-related PD neuropathology.

Multiple studies examining LRRK2 carriers (see Suppl. Table 2 for comprehensive list) have shown that LB-negative cases display Alzheimer’s disease (AD) or PSP-like tau pathology or show unspecific nigral cell loss without any underlying pathology [9, 32, 68, 82, 87, 90]. However, other pleiomorphic pathologies also include MSA and argyrophilic grain disease [31, 61]. Except for the PSP-diagnosed LRRK2 4, which was confirmed by neuropathology, tau pathology was not prominent among the LRRK2 cases in this cohort, and we found no correlation of particulate PLA with either Braak tangle stage or Aβ Thal phase. The sparse tau pathology in our LRRK2 cases is somewhat contrasting with the findings from Henderson et al. who examined a cohort of 11 LRRK2 cases, of which four were LB-negative [32]. These cases, mostly with G2019S mutation, all contained AD-like tau pathology, though levels in LB-negative cases tended to be lower than in the LB-positive cases [32]. Whether the divergent neuropathological findings are related to specific LRRK2 mutations is yet to be determined in larger cohort studies, as is the exact relationship between particulate PLA and other protein pathologies.

For most of the healthy control cases, stained in parallel with LRRK2 and regular LBD cases, only very low levels of particulate PLA were detected. None of the healthy controls showed any LBs and only had limited, if any, other neurodegenerative proteinaceous inclusions, such as Aβ plaques and tau tangles. Two of the controls (Control 2 and 6), however, did present with higher levels of particulate PLA, which were closer to the levels seen in some of the LBD and LRRK2 cases. This was seen particularly in the amygdala and SNpc but not in DMV or LC, which are usually thought to be affected early on with Lewy-like pathology. Previous studies using α-synuclein PLA have shown that it appears to detect earlier aggregate species than the Lewy pathology [36, 63, 76]. These two controls could thus represent early asymptomatic stages of LBD, perhaps of the brain-first subtype [10, 33, 34], that would have become symptomatic, had they lived longer. As we did have two other controls (Control 4 and 5) that were virtually void of any PLA signal in the study, it is unlikely that the α-synuclein PLA accumulation is a normal, disease-unrelated process. Whether there is a certain threshold above which the PLA-positive species become problematic and how they progress in the brain is currently unknown and requires further studies in larger cohorts of early-stage LBD cases and aged controls.

The main limitation of this study pertains to the relatively small cohort size, which relates to the scarcity of LB-negative LRRK2 cases available for post-mortem examination. As such, larger studies are needed to determine whether the non-inclusion α-synuclein aggregate pathology is a general feature of LRRK2 cases and how its extent and distribution may correlate with other protein pathologies. Additionally, the inclusion of cases with different clinical diagnoses (within both LRRK2 and LBD groups) could obscure features that are specific for one clinical diagnosis but were not apparent in our study due to group heterogeneity. The study is also limited by the lack of complementary methods to confirm the presence of non-inclusion α-synuclein aggregates in the LRRK2 cases without Lewy pathology. One strategy for future studies could be to perform SAA on brain homogenates from regions with and without particulate PLA signal, which would inform on the presence of seeding-competent α-synuclein species in these regions. Nevertheless, we here provide compelling evidence that Lewy pathology is not representative of all α-synuclein aggregates in the brain.

In conclusion, we demonstrate substantial levels of α-synuclein aggregates, but no Lewy-like inclusions, in LB-negative LRRK2 cases. The particulate PLA affects both classical PD-related brain regions in the LRRK2 cases, but also other regions not usually implicated in PD, particularly the inferior olivary nucleus and the pontocerebellar pathways. These results motivate further studies on not only LRRK2 cases but also PRKN PD cases, which also frequently present without LBs upon autopsy. Furthermore, we need to also examine larger aged cohorts to understand how this particulate PLA signal progresses in the brain and when it becomes associated to clinical symptoms.

## Data availability

All data are presented in the manuscript or the supplementary material. Raw data supporting the findings of this study are available from the corresponding author upon reasonable request.

## Acknowledgements

The authors would like to sincerely thank all the brain donors for their contribution to this research. We acknowledge the Oxford Brain Bank, supported by the Medical Research Council (MRC), Brains for Dementia Research (BDR) (Alzheimer Society and Alzheimer Research UK), and the NIHR Oxford Biomedical Research Centre, as well as Imperial Brain Bank supported by Parkinson’s UK for supplying tissue. We further acknowledge the Imaging Core at the Department of Biomedicine, Aarhus University, for the use of facilities and assistance in imaging.

## Author information

### Author affiliations

DANDRITE - Danish Research Institute of Translational Neuroscience, Aarhus C, Denmark. Nanna Møller Jensen, Zagorka Vitic, Mia R. Antorini, Tobias Bruun Viftrup, Poul Henning Jensen.

Department of Biomedicine, Aarhus University, Aarhus C, Denmark.

Nanna Møller Jensen, Zagorka Vitic, Mia R. Antorini, Tobias Bruun Viftrup, Poul Henning Jensen.

Oxford Parkinson’s Disease Centre, University of Oxford, Oxford, United Kingdom. Laura Parkkinen.

Nuffield Department of Clinical Neurosciences, University of Oxford, Oxford, United Kingdom. Laura Parkkinen.

### Author contributions

NMJ, LP, and PHJ conceived and designed the study, and LP obtained post-mortem brain samples. NMJ, ZV, MRA, and TBV performed the experiments and analysed the data, while LP and PHJ provided critical input for the interpretation of data. Figures and first draft of the manuscript were prepared by NMJ. All authors provided feedback and approved the final version of the manuscript.

## Ethics declarations

### Ethical approval and informed consent

Post-mortem brain tissue was obtained from Parkinson’s UK Brain Bank (PUKBB), Imperial College London, and Oxford Brain Bank (OBB), Nuffield Department of Clinical Neurosciences in University of Oxford, in accordance with approved protocols by the Wales Research Ethics Committee (23/WA/0273) and the Ethics Committee of the University of Oxford (ref 23/SC/0241). All participants had given prior written informed consent for the brain donation. Both brain banks comply with the requirements of the Human Tissue Act 2004 and the Codes of Practice set by the Human Tissue Authority (HTA licence numbers 12275 for PUKBB and 12217 for OBB).

## References

1. Altay MF, Liu AKL, Holton JL, Parkkinen L, Lashuel HA (2022) Prominent astrocytic alpha-synuclein pathology with unique post-translational modification signatures unveiled across Lewy body disorders. Acta Neuropathol Commun 10:1–18. doi: 10.1186/s40478-022-01468-8

2. Amunts K, Kedo O, Kindler M, Pieperhoff P, Mohlberg H, Shah NJ, Habel U, Schneider F, Zilles K (2005) Cytoarchitectonic mapping of the human amygdala, hippocampal region and entorhinal cortex: Intersubject variability and probability maps. Anat Embryol (Berl) 210:343–352. doi: 10.1007/S00429-005-0025-5

3. Bankhead P, Loughrey MB, Fernández JA, Dombrowski Y, McArt DG, Dunne PD, McQuaid S, Gray RT, Murray LJ, Coleman HG, James JA, Salto-Tellez M, Hamilton PW (2017) QuPath: Open source software for digital pathology image analysis. Scientific Reports 2017 7:1 7:1–7. doi: 10.1038/s41598-017-17204-5

4. Beach TG, Adler CH, Lue LF, Sue LI, Bachalakuri J, Henry-Watson J, Sasse J, Boyer S, Shirohi S, Brooks R, Eschbacher J, White CL, Akiyama H, Caviness J, Shill HA, Connor DJ, Sabbagh MN, Walker DG (2009) Unified staging system for Lewy body disorders: Correlation with nigrostriatal degeneration, cognitive impairment and motor dysfunction. Acta Neuropathol 117:613–634. doi: 10.1007/s00401-009-0538-8

5. Bengoa-Vergniory N, Roberts RF, Wade-Martins R, Alegre-Abarrategui J (2017) Alpha-synuclein oligomers: a new hope. Acta Neuropathol 134:819–838. doi: 10.1007/s00401-017-1755-1

6. Bentivenga GM, Mammana A, Baiardi S, Rossi M, Ticca A, Magliocchetti F, Mastrangelo A, Poleggi A, Ladogana A, Capellari S, Parchi P (2024) Performance of a seed amplification assay for misfolded alpha-synuclein in cerebrospinal fluid and brain tissue in relation to Lewy body disease stage and pathology burden. Acta Neuropathol 147:1–13. doi: 10.1007/S00401-023-02663-0

7. Berkhoudt Lassen L, Gregersen E, Kathrine Isager A, Betzer C, Hahn Kofoed R, Henning Jensen P (2018) ELISA method to detect α-synuclein oligomers in cell and animal models. PLoS One 13:e0196056. doi: 10.1371/JOURNAL.PONE.0196056

8. Berwick DC, Heaton GR, Azeggagh S, Harvey K (2019) LRRK2 Biology from structure to dysfunction: research progresses, but the themes remain the same. Molecular Neurodegeneration 2019 14:1 14:1–22. doi: 10.1186/S13024-019-0344-2

9. Blauwendraat C, Pletnikova O, Geiger JT, Murphy NA, Abramzon Y, Rudow G, Mamais A, Sabir MS, Crain B, Ahmed S, Rosenthal LS, Bakker CC, Faghri F, Chia R, Ding J, Dawson TM, Pantelyat A, Albert MS, Nalls MA, Resnick SM, Ferrucci L, Cookson MR, Hillis AE, Troncoso JC, Scholz SW (2019) Genetic analysis of neurodegenerative diseases in a pathology cohort. Neurobiol Aging 76:214.e1–214.e9. doi: 10.1016/J.NEUROBIOLAGING.2018.11.007

10. Borghammer P, Horsager J, Andersen K, Van Den Berge N, Raunio A, Murayama S, Parkkinen L, Myllykangas L (2021) Neuropathological evidence of body-first vs. brain-first Lewy body disease. Neurobiol Dis 161:105557. doi: 10.1016/j.nbd.2021.105557

11. Braak H, Ghebremedhin E, Rüb U, Bratzke H, Del Tredici K (2004) Stages in the development of Parkinson’s disease-related pathology. Cell Tissue Res 318:121–134

12. Braak H, Del Tredici K, Rüb U, De Vos RAI, Jansen Steur ENH, Braak E (2003) Staging of brain pathology related to sporadic Parkinson’s disease. Neurobiol Aging 24:197–211. doi: 10.1016/S0197-4580(02)00065-9

13. Cabezudo D, Tsafaras G, Van Acker E, Van den Haute C, Baekelandt V (2023) Mutant LRRK2 exacerbates immune response and neurodegeneration in a chronic model of experimental colitis. Acta Neuropathol 146:245–261. doi: 10.1007/S00401-023-02595-9

14. Cherian A, Divya KP (2020) Genetics of Parkinson’s disease. Acta Neurol Belg 120:1297–1305. doi: 10.1007/S13760-020-01473-5

15. Covy JP, Yuan W, Waxman EA, Hurtig HI, Van Deerlin VM, Giasson BI (2009) Clinical and pathological characteristics of patients with Leucine-rich repeat kinase-2 mutations. Movement Disorders 24:32–39. doi: 10.1002/MDS.22096

16. Dächsel JC, Ross OA, Mata IF, Kachergus J, Toft M, Cannon A, Baker M, Adamson J, Hutton M, Dickson DW, Farrer MJ (2007) Lrrk2 G2019S substitution in frontotemporal lobar degeneration with ubiquitin-immunoreactive neuronal inclusions. Acta Neuropathol 113:601–606. doi: 10.1007/S00401-006-0178-1

17. Day JO, Mullin S (2021) The Genetics of Parkinson’s Disease and Implications for Clinical Practice. Genes 2021, Vol 12, Page 1006 12:1006. doi: 10.3390/GENES12071006

18. Dickson DW, Braak H, Duda JE, Duyckaerts C, Gasser T, Halliday GM, Hardy J, Leverenz JB, Del Tredici K, Wszolek ZK, Litvan I (2009) Neuropathological assessment of Parkinson’s disease: refining the diagnostic criteria. Lancet Neurol 8:1150–1157

19. Doherty KM, Silveira-Moriyama L, Parkkinen L, Healy DG, Farrell M, Mencacci NE, Ahmed Z, Brett FM, Hardy J, Quinn N, Counihan TJ, Lynch T, Fox Z V., Revesz T, Lees AJ, Holton JL (2013) Parkin Disease: A Clinicopathologic Entity? JAMA Neurol 70:571–579. doi: 10.1001/JAMANEUROL.2013.172

20. Duffy PE, Tennyson VM (1965) Phase and Electron Microscopic Observations of Lewy Bodies and Melanin Granules in the Substantia Nigra and Locus Caeruleus in Parkinson’s Disease. J Neuropathol Exp Neurol 24:398–414. doi: 10.1097/00005072-196507000-00003

21. Gai WP, Blumbergs PC, Geffen LB, Blessing WW (1992) Age-related loss of dorsal vagal neurons in Parkinson’s disease. Neurology 42:2106–2111. doi: 10.1212/WNL.42.11.2106

22. Gaig C, Martí MJ, Ezquerra M, Rey MJ, Cardozo A, Tolosa E (2007) G2019S LRRK2 mutation causing Parkinson’s disease without Lewy bodies. J Neurol Neurosurg Psychiatry 78:626–628. doi: 10.1136/JNNP.2006.107904

23. German DC, Manaye KF, White CL, Woodward DJ, McIntire DD, Smith WK, Kalaria RN, Mann DMA (1992) Disease-specific patterns of locus coeruleus cell loss. Ann Neurol 32:667–676. doi: 10.1002/ANA.410320510

24. Giasson BI, Covy JP, Bonini NM, Hurtig HI, Farrer MJ, Trojanowski JQ, Van Deerlin VM (2006) Biochemical and pathological characterization of Lrrk2. Ann Neurol 59:315–322. doi: 10.1002/ANA.20791

25. Gilks WP, Abou-Sleiman PM, Gandhi S, Jain S, Singleton A, Lees AJ, Shaw K, Bhatia KP, Bonifati V, Quinn NP, Lynch J, Healy DG, Holton JL, Revesz T, Wood NW (2005) A common LRRK2 mutation in idiopathic Parkinson’s disease. The Lancet 365:415–416. doi: 10.1016/S0140-6736(05)17830-1

26. Giordana MT, D’Agostino C, Albani G, Mauro A, Di Fonzo A, Antonini A, Bonifati V (2007) Neuropathology of Parkinson’s disease associated with the LRRK2 Ile1371Val mutation. Movement Disorders 22:275–278. doi: 10.1002/MDS.21281

27. Gomez A, Ferrer I (2010) Involvement of the cerebral cortex in Parkinson disease linked with G2019S LRRK2 mutation without cognitive impairment. Acta Neuropathol 120:155–167. doi: 10.1007/S00401-010-0669-Y

28. Gómez-Suaga P, Luzón-Toro B, Churamani D, Zhang L, Bloor-Young D, Patel S, Woodman PG, Churchill GC, Hilfiker S (2012) Leucine-rich repeat kinase 2 regulates autophagy through a calcium-dependent pathway involving NAADP. Hum Mol Genet 21:511–525. doi: 10.1093/HMG/DDR481

29. Halliday G, Reyes S, Double K (2012) Substantia Nigra, Ventral Tegmental Area, and Retrorubral Fields. The Human Nervous System, Third Edition 439–455. doi: 10.1016/B978-0-12-374236-0.10013-6

30. Harding AJ, Halliday GM (2001) Cortical Lewy body pathology in the diagnosis of dementia. Acta Neuropathol 102:355–363. doi: 10.1007/S004010100390

31. Hasegawa K, Stoessl AJ, Yokoyama T, Kowa H, Wszolek ZK, Yagishita S (2009) Familial parkinsonism: study of original Sagamihara PARK8 (I2020T) kindred with variable clinicopathologic outcomes. Parkinsonism Relat Disord 15:300–306. doi: 10.1016/J.PARKRELDIS.2008.07.010

32. Henderson MX, Sengupta M, Trojanowski JQ, Lee VMY (2019) Alzheimer’s disease tau is a prominent pathology in LRRK2 Parkinson’s disease. Acta Neuropathol Commun 7:183. doi: 10.1186/s40478-019-0836-x

33. Horsager J, Andersen KB, Knudsen K, Skjærbæk C, Fedorova TD, Okkels N, Schaeffer E, Bonkat SK, Geday J, Otto M, Sommerauer M, Danielsen EH, Bech E, Kraft J, Munk OL, Hansen SD, Pavese N, Göder R, Brooks DJ, Berg D, Borghammer P (2020) Brain-first versus body-first Parkinson’s disease: A multimodal imaging case-control study. Brain 143:3077–3088. doi: 10.1093/brain/awaa238

34. Horsager J, Borghammer P (2024) Brain-first vs. body-first Parkinson’s disease: An update on recent evidence. Parkinsonism Relat Disord 122:106101

35. Hui KY, Fernandez-Hernandez H, Hu J, Schaffner A, Pankratz N, Hsu NY, Chuang LS, Carmi S, Villaverde N, Li X, Rivas M, Levine AP, Bao X, Labrias PR, Haritunians T, Ruane D, Gettler K, Chen E, Li D, Schiff ER, Pontikos N, Barzilai N, Brant SR, Bressman S, Cheifetz AS, Clark LN, Daly MJ, Desnick RJ, Duerr RH, Katz S, Lencz T, Myers RH, Ostrer H, Ozelius L, Payami H, Peter Y, Rioux JD, Segal AW, Scott WK, Silverberg MS, Vance JM, Ubarretxena-Belandia I, Foroud T, Atzmon G, Pe’er I, Ioannou Y, McGovern DPB, Yue Z, Schadt EE, Cho JH, Peter I (2018) Functional variants in the LRRK2 gene confer shared effects on risk for Crohn’s disease and Parkinson’s disease. Sci Transl Med 10:7795.

36. Jensen NM, Fu Y, Betzer C, Li H, Elfarrash S, Shaib AH, Krah D, Vitic Z, Reimer L, Gram H, Buchman V, Denham M, Rizzoli SO, Halliday GM, Jensen PH (2024) MJF-14 proximity ligation assay detects early non-inclusion alpha-synuclein pathology with enhanced specificity and sensitivity. NPJ Parkinsons Dis 10:227. doi: 10.1038/s41531-024-00841-9

37. Kalia L V, Lang AE (2015) Parkinson’s disease. The Lancet 386:896–912. doi: 10.1016/S0140-6736(14)61393-3

38. Kalia L V., Lang AE, Hazrati LN, Fujioka S, Wszolek ZK, Dickson DW, Ross OA, Van Deerlin VM, Trojanowski JQ, Hurtig HI, Alcalay RN, Marder KS, Clark LN, Gaig C, Tolosa E, Ruiz-Martínez J, Marti-Masso JF, Ferrer I, López De Munain A, Goldman SM, Schüle B, Langston JW, Aasly JO, Giordana MT, Bonifati V, Puschmann A, Canesi M, Pezzoli G, Maues De Paula A, Hasegawa K, Duyckaerts C, Brice A, Stoessl AJ, Marras C (2015) Clinical Correlations With Lewy Body Pathology in LRRK2-Related Parkinson Disease. JAMA Neurol 72:100–105. doi: 10.1001/JAMANEUROL.2014.2704

39. Kalogeropulou AF, Purlyte E, Tonelli F, Lange SM, Wightman M, Prescott AR, Padmanabhan S, Sammler E, Alessi DR (2022) Impact of 100 LRRK2 variants linked to Parkinson’s disease on kinase activity and microtubule binding. Biochemical Journal 479:1759–1783. doi: 10.1042/BCJ20220161

40. Khan NL, Jain S, Lynch JM, Pavese N, Abou-Sleiman P, Holton JL, Healy DG, Gilks WP, Sweeney MG, Ganguly M, Gibbons V, Gandhi S, Vaughan J, Eunson LH, Katzenschlager R, Gayton J, Lennox G, Revesz T, Nicholl D, Bhatia KP, Quinn N, Brooks D, Lees AJ, Davis MB, Piccini P, Singleton AB, Wood NW (2005) Mutations in the gene LRRK2 encoding dardarin (PARK8) cause familial Parkinson’s disease: clinical, pathological, olfactory and functional imaging and genetic data. Brain 128:2786–2796. doi: 10.1093/BRAIN/AWH667

41. Kramer ML, Schulz-Schaeffer WJ (2007) Presynaptic α-synuclein aggregates, not Lewy bodies, cause neurodegeneration in dementia with lewy bodies. Journal of Neuroscience 27:1405–1410. doi: 10.1523/JNEUROSCI.4564-06.2007

42. Lake J, Reed X, Langston RG, Nalls MA, Gan-Or Z, Cookson MR, Singleton AB, Blauwendraat C, Leonard HL, Agid Y, Anheim M, Artaud F, Bonnet AM, Bonnet C, Bourdain F, Brandel JP, Brefel-Courbon C, Borg M, Brice A, Broussolle E, Cormier-Dequaire F, Corvol JC, Damier P, Debilly B, Degos B, Derkinderen P, Destée A, Dürr A, Durif F, Elbaz A, Grabli D, Hartmann A, Klebe S, Krack P, Kraemmer J, Leder S, Lesage S, Levy R, Lohmann E, Lacomblez L, Mangone G, Mariani LL, Marques AR, Martinez M, Mesnage V, Muellner J, Ory-Magne F, Pico F, Planté-Bordeneuve V, Pollak P, Rascol O, Tahiri K, Tison F, Tranchant C, Roze E, Tir M, Vérin M, Viallet F, Vidailhet M, You A (2022) Coding and Noncoding Variation in LRRK2 and Parkinson’s Disease Risk. Movement Disorders 37:95–105. doi: 10.1002/MDS.28787

43. Lashuel HA (2020) Do Lewy bodies contain alpha-synuclein fibrils? and Does it matter? A brief history and critical analysis of recent reports. Neurobiol Dis 141:104876

44. Ling H, Kara E, Bandopadhyay R, Hardy J, Holton J, Xiromerisiou G, Lees A, Houlden H, Revesz T (2013) TDP-43 pathology in a patient carrying G2019S LRRK2 mutation and a novel p.Q124E MAPT. Neurobiol Aging 34:2889.e5-2889.e9. doi: 10.1016/J.NEUROBIOLAGING.2013.04.011

45. Madureira M, Connor-Robson N, Wade-Martins R (2020) “LRRK2: Autophagy and Lysosomal Activity.” Front Neurosci 14:536324. doi: 10.3389/FNINS.2020.00498/BIBTEX

46. Mahul-Mellier AL, Burtscher J, Maharjan N, Weerens L, Croisier M, Kuttler F, Leleu M, Knott GW, Lashuel HA (2020) The process of Lewy body formation, rather than simply α-synuclein fibrillization, is one of the major drivers of neurodegeneration. Proc Natl Acad Sci U S A 117:4971–4982. doi: 10.1073/pnas.1913904117

47. Mahul-Mellier AL, Vercruysse F, MacO B, Ait-Bouziad N, De Roo M, Muller D, Lashuel HA (2015) Fibril growth and seeding capacity play key roles in α-synuclein-mediated apoptotic cell death. Cell Death Differ 22:2107–2122. doi: 10.1038/cdd.2015.79

48. Martí-Massó JF, Ruiz-Martínez J, Bolaño MJ, Ruiz I, Gorostidi A, Moreno F, Ferrer I, López De Munain A (2009) Neuropathology of Parkinson’s disease with the R1441G mutation in LRRK2. Mov Disord 24:1998–2001. doi: 10.1002/MDS.22677

49. McKeith IG, Boeve BF, Dickson DW, Halliday G, Taylor JP, Weintraub D, Aarsland D, Galvin J, Attems J, Ballard CG, Bayston A, Beach TG, Blanc F, Bohnen N, Bonanni L, Bras J, Brundin P, Burn D, Chen-Plotkin A, Duda JE, El-Agnaf O, Feldman H, Ferman TJ, Ffytche D, Fujishiro H, Galasko D, Goldman JG, Gomperts SN, Graff-Radford NR, Honig LS, Iranzo A, Kantarci K, Kaufer D, Kukull W, Lee VMY, Leverenz JB, Lewis S, Lippa C, Lunde A, Masellis M, Masliah E, McLean P, Mollenhauer B, Montine TJ, Moreno E, Mori E, Murray M, O’Brien JT, Orimo S, Postuma RB, Ramaswamy S, Ross OA, Salmon DP, Singleton A, Taylor A, Thomas A, Tiraboschi P, Toledo JB, Trojanowski JQ, Tsuang D, Walker Z, Yamada M, Kosaka K (2017) Diagnosis and management of dementia with Lewy bodies. Neurology 89:88–100

50. Mulroy E, Erro R, Bhatia KP, Hallett M (2024) Refining the clinical diagnosis of Parkinson’s disease. Parkinsonism Relat Disord 122

51. Nakano I, Hirano A (1984) Parkinson’s disease: neuron loss in the nucleus basalis without concomitant Alzheimer’s disease. Ann Neurol 15:415–418. doi: 10.1002/ANA.410150503

52. Nalls MA, Bras J, Hernandez DG, Keller MF, Majounie E, Renton AE, Saad M, Jansen I, Guerreiro R, Lubbe S, Plagnol V, Gibbs JR, Schulte C, Pankratz N, Sutherland M, Bertram L, Lill CM, Destefano AL, Faroud T, Eriksson N, Tung JY, Edsall C, Nichols N, Brooks J, Arepalli S, Pliner H, Letson C, Heutink P, Martinez M, Gasser T, Traynor BJ, Wood N, Hardy J, Singleton AB, Agid Y, Anheim M, Bonnet AM, Borg M, Brice A, Broussolle E, Corvol JC, Damier P, Destée A, Dürr A, Durif F, Klebe S, Lohmann E, Martinez M, Pollak P, Rascol O, Tison F, Tranchant C, Vérin M, Viallet F, Vidailhet M, Alpérovitch A, Berr C, Tzourio C (2015) NeuroX, a fast and efficient genotyping platform for investigation of neurodegenerative diseases. Neurobiol Aging 36:1605.e7–1605.e12. doi: 10.1016/J.NEUROBIOLAGING.2014.07.028

53. Nalls MA, Pankratz N, Lill CM, Do CB, Hernandez DG, Saad M, DeStefano AL, Kara E, Bras J, Sharma M, Schulte C, Keller MF, Arepalli S, Letson C, Edsall C, Stefansson H, Liu X, Pliner H, Lee JH, Cheng R, Ikram MA, Ioannidis JPA, Hadjigeorgiou GM, Bis JC, Martinez M, Perlmutter JS, Goate A, Marder K, Fiske B, Sutherland M, Xiromerisiou G, Myers RH, Clark LN, Stefansson K, Hardy JA, Heutink P, Chen H, Wood NW, Houlden H, Payami H, Brice A, Scott WK, Gasser T, Bertram L, Eriksson N, Foroud T, Singleton AB (2014) Large-scale meta-analysis of genome-wide association data identifies six new risk loci for Parkinson’s disease. Nat Genet 46:989–993. doi: 10.1038/ng.3043

54. Neumann M, Kahle PJ, Giasson BI, Ozmen L, Borroni E, Spooren W, Müller V, Odoy S, Fujiwara H, Hasegawa M, Iwatsubo T, Trojanowski JQ, Kretzschmar HA, Haass C (2002) Misfolded proteinase K–resistant hyperphosphorylated α-synuclein in aged transgenic mice with locomotor deterioration and in human α-synucleinopathies. Journal of Clinical Investigation 110:1429–1439. doi: 10.1172/JCI15777

55. Olanow CW, Perl DP, DeMartino GN, McNaught KSP (2004) Lewy-body formation is an aggresome-related process: A hypothesis. Lancet Neurology 3:496–503

56. Outeiro TF, Koss DJ, Erskine D, Walker L, Kurzawa-Akanbi M, Burn D, Donaghy P, Morris C, Taylor JP, Thomas A, Attems J, McKeith I (2019) Dementia with Lewy bodies: An update and outlook. Mol Neurodegener 14:1–18

57. Pieri L, Madiona K, Melki R (2016) Structural and functional properties of prefibrillar α-synuclein oligomers. Sci Rep 6:1–15. doi: 10.1038/srep24526

58. Polymeropoulos MH (1997) Mutation in the α-Synuclein Gene Identified in Families with Parkinson’s Disease. Science (1979) 276:2045–2047. doi: 10.1126/science.276.5321.2045

59. Poulopoulos M, Levy OA, Alcalay RN (2012) The neuropathology of genetic Parkinson’s disease. Movement Disorders 27:831–842. doi: 10.1002/MDS.24962

60. Puschmann A, Englund E, Ross OA, Vilariño-Güell C, Lincoln SJ, Kachergus JM, Cobb SA, Törnqvist AL, Rehncrona S, Widner H, Wszolek ZK, Farrer MJ, Nilsson C (2012) First neuropathological description of a patient with Parkinson’s disease and LRRK2 p.N1437H mutation. Parkinsonism Relat Disord 18:332–338. doi: 10.1016/J.PARKRELDIS.2011.11.019

61. Rajput A, Dickson DW, Robinson CA, Ross OA, Dächsel JC, Lincoln SJ, Cobb SA, Rajput ML, Farrer MJ (2006) Parkinsonism, Lrrk2 G2019S, and tau neuropathology. Neurology 67:1506–1508. doi: 10.1212/01.WNL.0000240220.33950.0C

62. Refai FS, Ng SH, Tan EK (2015) Evaluating LRRK2 Genetic Variants with Unclear Pathogenicity. Biomed Res Int 2015:678701. doi: 10.1155/2015/678701

63. Roberts RF, Wade-Martins R, Alegre-Abarrategui J (2015) Direct visualization of alpha-synuclein oligomers reveals previously undetected pathology in Parkinson’s disease brain. Brain 138:1642–1657. doi: 10.1093/brain/awv040

64. Ross OA, Soto-Ortolaza AI, Heckman MG, Aasly JO, Abahuni N, Annesi G, Bacon JA, Bardien S, Bozi M, Brice A, Brighina L, Van Broeckhoven C, Carr J, Chartier-Harlin MC, Dardiotis E, Dickson DW, Diehl NN, Elbaz A, Ferrarese C, Ferraris A, Fiske B, Gibson JM, Gibson R, Hadjigeorgiou GM, Hattori N, Ioannidis JPA, Jasinska-Myga B, Jeon BS, Kim YJ, Klein C, Kruger R, Kyratzi E, Lesage S, Lin CH, Lynch T, Maraganore DM, Mellick GD, Mutez E, Nilsson C, Opala G, Park SS, Puschmann A, Quattrone A, Sharma M, Silburn PA, Sohn YH, Stefanis L, Tadic V, Theuns J, Tomiyama H, Uitti RJ, Valente EM, van de Loo S, Vassilatis DK, Vilariño-Güell C, White LR, Wirdefeldt K, Wszolek ZK, Wu RM, Farrer MJ (2011) Association of LRRK2 exonic variants with susceptibility to Parkinson’s disease: a case–control study. Lancet Neurol 10:898–908. doi: 10.1016/S1474-4422(11)70175-2

65. Ross OA, Toft M, Whittle AJ, Johnson JL, Papapetropoulos S, Mash DC, Litvan I, Gordon MF, Wszolek ZK, Farrer MJ, Dickson DW (2006) Lrrk2 and Lewy body disease. Ann Neurol 59:388–393. doi: 10.1002/ANA.20731

66. Ruesink H, Reimer L, Gregersen E, Moeller A, Betzer C, Jensen PH (2019) Stabilization of α-synuclein oligomers using formaldehyde. PLoS One 14:e0216764. doi: 10.1371/journal.pone.0216764

67. Ruffmann C, Giaccone G, Canesi M, Bramerio M, Goldwurm S, Gambacorta M, Rossi G, Tagliavini F, Pezzoli G (2012) Atypical tauopathy in a patient with LRRK2-G2019S mutation and tremor-dominant Parkinsonism. Neuropathol Appl Neurobiol 38:382–386. doi: 10.1111/J.1365-2990.2011.01216.X

68. Sanchez-Contreras M, Heckman MG, Tacik P, Diehl N, Brown PH, Soto-Ortolaza AI, Christopher EA, Walton RL, Ross OA, Golbe LI, Graff-Radford N, Wszolek ZK, Dickson DW, Rademakers R (2017) Study of LRRK2 variation in tauopathy: Progressive supranuclear palsy and corticobasal degeneration. Movement Disorders 32:115–123. doi: 10.1002/MDS.26815

69. Schapansky J, Khasnavis S, DeAndrade MP, Nardozzi JD, Falkson SR, Boyd JD, Sanderson JB, Bartels T, Melrose HL, LaVoie MJ (2018) Familial knockin mutation of LRRK2 causes lysosomal dysfunction and accumulation of endogenous insoluble α-synuclein in neurons. Neurobiol Dis 111:26–35. doi: 10.1016/J.NBD.2017.12.005

70. Schapira AHV, Chaudhuri KR, Jenner P (2017) Non-motor features of Parkinson disease. Nature Reviews Neuroscience 2017 18:7 18:435–450. doi: 10.1038/nrn.2017.62

71. Schneider SA, Alcalay RN (2017) Neuropathology of genetic synucleinopathies with parkinsonism: Review of the literature. Movement Disorders 32:1504–1523. doi: 10.1002/MDS.27193

72. Schröder H, de Vos RAI, Huggenberger S, Müller-Thomsen L, Rozemuller A, Hedayat F, Moser N (2023) Histology atlas of the human brainstem. In: The Human Brainstem. Springer, Cham, pp 539–630

73. Schulz-Schaeffer WJ (2010) The synaptic pathology of α-synuclein aggregation in dementia with Lewy bodies, Parkinson’s disease and Parkinson’s disease dementia. Acta Neuropathol 120:131–143. doi: 10.1007/S00401-010-0711-0

74. Seidel K, Mahlke J, Siswanto S, Krüger R, Heinsen H, Auburger G, Bouzrou M, Grinberg LT, Wicht H, Korf HW, Den Dunnen W, Rüb U (2015) The Brainstem Pathologies of Parkinson’s Disease and Dementia with Lewy Bodies. Brain Pathology 25:121–135. doi: 10.1111/BPA.12168

75. Sekiya H, Kowa H, Koga H, Takata M, Satake W, Futamura N, Funakawa I, Jinnai K, Takahashi M, Kondo T, Ueno Y, Kanagawa M, Kobayashi K, Toda T (2019) Wide distribution of alpha-synuclein oligomers in multiple system atrophy brain detected by proximity ligation. Acta Neuropathol 137:455–466. doi: 10.1007/s00401-019-01961-w

76. Sekiya H, Tsuji A, Hashimoto Y, Takata M, Koga S, Nishida K, Futamura N, Kawamoto M, Kohara N, Dickson DW, Kowa H, Toda T (2022) Discrepancy between distribution of alpha-synuclein oligomers and Lewy-related pathology in Parkinson’s disease. Acta Neuropathol Commun 10:133. doi: 10.1186/s40478-022-01440-6

77. Shahmoradian SH, Lewis AJ, Genoud C, Hench J, Moors TE, Navarro PP, Castaño-Díez D, Schweighauser G, Graff-Meyer A, Goldie KN, Sütterlin R, Huisman E, Ingrassia A, Gier Y de, Rozemuller AJM, Wang J, Paepe A De, Erny J, Staempfli A, Hoernschemeyer J, Großerüschkamp F, Niedieker D, El-Mashtoly SF, Quadri M, Van IJcken WFJ, Bonifati V, Gerwert K, Bohrmann B, Frank S, Britschgi M, Stahlberg H, Van de Berg WDJ, Lauer ME (2019) Lewy pathology in Parkinson’s disease consists of crowded organelles and lipid membranes. Nature Neuroscience 2019 22:7 22:1099–1109. doi: 10.1038/s41593-019-0423-2

78. Siderowf A, Concha-Marambio L, Lafontant DE, Farris CM, Ma Y, Urenia PA, Nguyen H, Alcalay RN, Chahine LM, Foroud T, Galasko D, Kieburtz K, Merchant K, Mollenhauer B, Poston KL, Seibyl J, Simuni T, Tanner CM, Weintraub D, Videnovic A, Choi SH, Kurth R, Caspell-Garcia C, Coffey CS, Frasier M, Oliveira LMA, Hutten SJ, Sherer T, Marek K, Soto C (2023) Assessment of heterogeneity among participants in the Parkinson’s Progression Markers Initiative cohort using α-synuclein seed amplification: a cross-sectional study. Lancet Neurol 22:407–417. doi: 10.1016/S1474-4422(23)00109-6

79. Sosero YL, Yu E, Krohn L, Rudakou U, Mufti K, Ruskey JA, Asayesh F, Laurent SB, Spiegelman D, Fahn S, Waters C, Sardi SP, Bandres-Ciga S, Alcalay RN, Gan-Or Z, Senkevich K (2021) LRRK2 p.M1646T is associated with glucocerebrosidase activity and with Parkinson’s disease. Neurobiol Aging 103: 142.e1–142.e5. doi: 10.1016/J.NEUROBIOLAGING.2021.02.018

80. Tanji K, Mori F, Mimura J, Itoh K, Kakita A, Takahashi H, Wakabayashi K (2010) Proteinase K-resistant α-synuclein is deposited in presynapses in human Lewy body disease and A53T α-synuclein transgenic mice. Acta Neuropathol 120:145–154. doi: 10.1007/s00401-010-0676-z

81. Trinkaus VA, Riera-Tur I, Martínez-Sánchez A, Bäuerlein FJB, Guo Q, Arzberger T, Baumeister W, Dudanova I, Hipp MS, Hartl FU, Fernández-Busnadiego R (2021) In situ architecture of neuronal α-Synuclein inclusions. Nature Communications 2021 12:1 12:1–10. doi: 10.1038/s41467-021-22108-0

82. Ujiie S, Hatano T, Kubo SI, Imai S, Sato S, Uchihara T, Yagishita S, Hasegawa K, Kowa H, Sakai F, Hattori N (2012) LRRK2 I2020T mutation is associated with tau pathology. Parkinsonism Relat Disord 18:819–823. doi: 10.1016/J.PARKRELDIS.2012.03.024

83. Vaikath NN, Majbour NK, Paleologou KE, Ardah MT, van Dam E, van de Berg WDJ, Forrest SL, Parkkinen L, Gai WP, Hattori N, Takanashi M, Lee SJ, Mann DMA, Imai Y, Halliday GM, Li JY, El-Agnaf OMA (2015) Generation and characterization of novel conformation-specific monoclonal antibodies for α-synuclein pathology. Neurobiol Dis 79:81–99. doi: 10.1016/J.NBD.2015.04.009

84. Wakabayashi K, Tanji K, Mori F, Takahashi H (2007) The Lewy body in Parkinson’s disease: Molecules implicated in the formation and degradation of α-synuclein aggregates. In: Neuropathology. John Wiley & Sons, Ltd (10.1111), pp 494–506

85. Wenning GK, Tison F, Elliott L, Quinn NP, Daniel SE (1996) Olivopontocerebellar pathology in multiple system atrophy. Mov Disord 11:157–162. doi: 10.1002/MDS.870110207

86. Woolsey TA, Hanaway J, Gado MH (2017) The Brain Atlas: A Visual Guide to the Human Central Nervous System, 3rd Edition, 4th ed

87. Wszolek ZK, Pfeiffer RF, Tsuboi Y, Uitti RJ, McComb RD, Stoessl AJ, Strongosky AJ, Zimprich A, Müller-Myhsok B, Farrer MJ, Gasser T, Calne DB, Dickson DW (2004) Autosomal dominant parkinsonism associated with variable synuclein and tau pathology. Neurology 62:1619–1622. doi: 10.1212/01.WNL.0000125015.06989.DB

88. Wszolek ZK, Vieregge P, Uitti RJ, Gasser T, Yasuhara O, McGeer P, Berry K, Calne DB, Vingerhoets FJG, Klein C, Pfeiffer RF (1997) German-Canadian family (Family A) with Parkinsonism, Amyotrophy, and Dementia-longitudinal observations. Parkinsonism Relat Disord 3:125–139. doi: 10.1016/S1353-8020(97)00013-8

89. Yilmazer-Hanke DM, Wolf HK, Schramm J, Elger CE, Wiestler OD, Blümcke I (2000) Subregional Pathology of the Amygdala Complex and Entorhinal Region in Surgical Specimens From Patients With Pharmacoresistant Temporal Lobe Epilepsy. J Neuropathol Exp Neurol 59

90. Zimprich A, Biskup S, Leitner P, Lichtner P, Farrer M, Lincoln S, Kachergus J, Hulihan M, Uitti RJ, Calne DB, Stoessl AJ, Pfeiffer RF, Patenge N, Carbajal IC, Vieregge P, Asmus F, Müller-Myhsok B, Dickson DW, Meitinger T, Strom TM, Wszolek ZK, Gasser T (2004) Mutations in LRRK2 Cause Autosomal-Dominant Parkinsonism with Pleomorphic Pathology. Neuron 44:601–607. doi: 10.1016/J.NEURON.2004.11.005

