## Supplementary material for "Abundant non-inclusion α-synuclein pathology in Lewy body-negative LRRK2-mutant cases"

Acta Neuropathologica (2025).

**Index**

| | |
|---|---|
| Suppl. Fig. 1 | p. 2 |
| Suppl. Fig. 2 | p. 3 |
| Suppl. Fig. 3 | p. 5 |
| Suppl. Fig. 4 | p. 6 |
| Suppl. Fig. 5 | p. 7 |
| Suppl. Table 1 | p. 8 |
| Suppl. Table 2 | p. 9 |

**Suppl. Fig. 1**

### Image analysis strategy

**1. Original image/ROI**

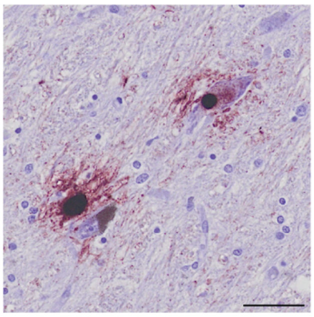

Original image to be analyzed (scale bar = 50 μm). Contains multiple types of signals, which need separation:

1. Neuromelanin (excluded from analysis)
2. PLA signal (particulate and Lewy-like subtypes)
3. Background tissue incl. neuronal and glial nuclei

**2. Neuromelanin definition (manual)**

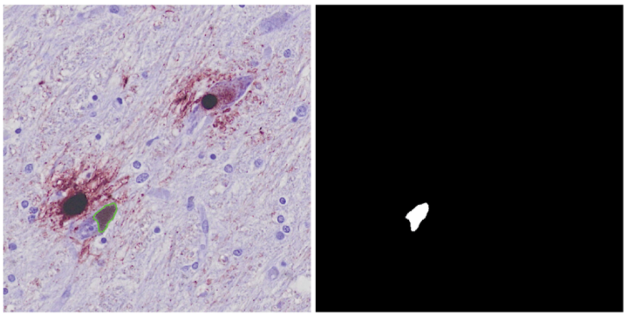

Neuronmelanin is manually outlined and excluded from further analysis (to avoid the algorithm mistaking neuro-melanin for PLA signal).

In the sample image, neuromelanin is outlined in green (left), while masked neuromelanin image is shown on the right.

**3. Total PLA definition (DAB thresholder)**

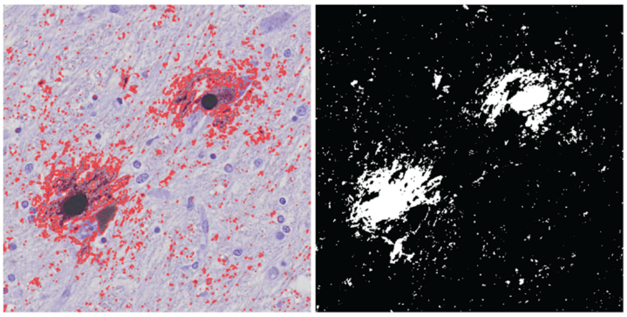

Total PLA signals (incl. both particulate and Lewy-like PLA) are defined based on thresholding on the DAB channel.
Thresholder criteria: Gaussian smoothing of 0.5, followed by a threshold of 0.1. Anything above the threshold is considered PLA signal.

In the sample image, total PLA is outlined in red (left), while masked total PLA image is shown on the right.

**4. Lewy-like PLA definition (average channel thresholder)**

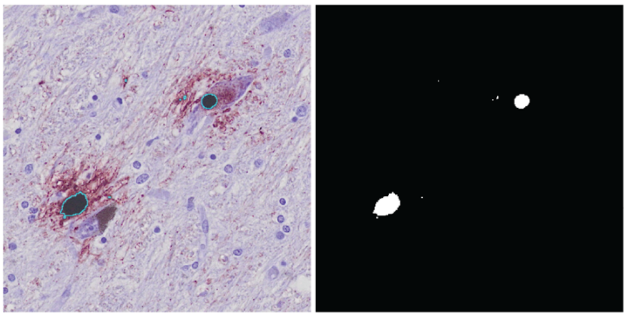

Lewy-like PLA is defined based on thresholding on the average channel intensity.
Thresholder criteria: Gaussian smoothing of 1.0, followed by a threshold of 80. Anything below the threshold is considered Lewy-like PLA.
Positive signal area is normalized to tissue area to create an area % measure.

In the sample image, Lewy-like PLA is outlined in cyan (left), while masked image is shown on the right.

**5. Particulate PLA (DAB – average channel thresholder)**

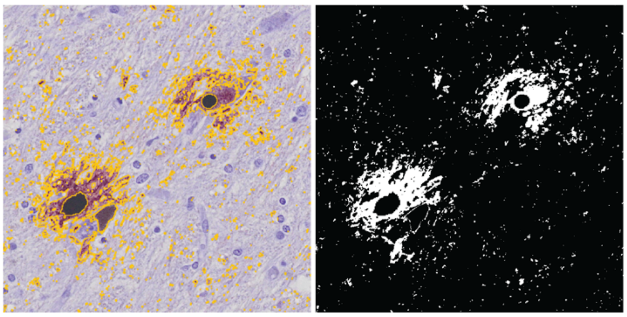

Particulate PLA is defined as the difference between the DAB thresholder (total PLA) and the average channel thresholder (Lewy-like PLA). I.e., particulate PLA is positive on the total PLA thresholder but negative on the Lewy-like PLA thresholder.
Positive signal area is normalized to tissue area to create an area % measure.

In the sample image, particulate PLA is outlined in yellow (left), while masked image is shown on the right.

**Suppl. Fig. 2**

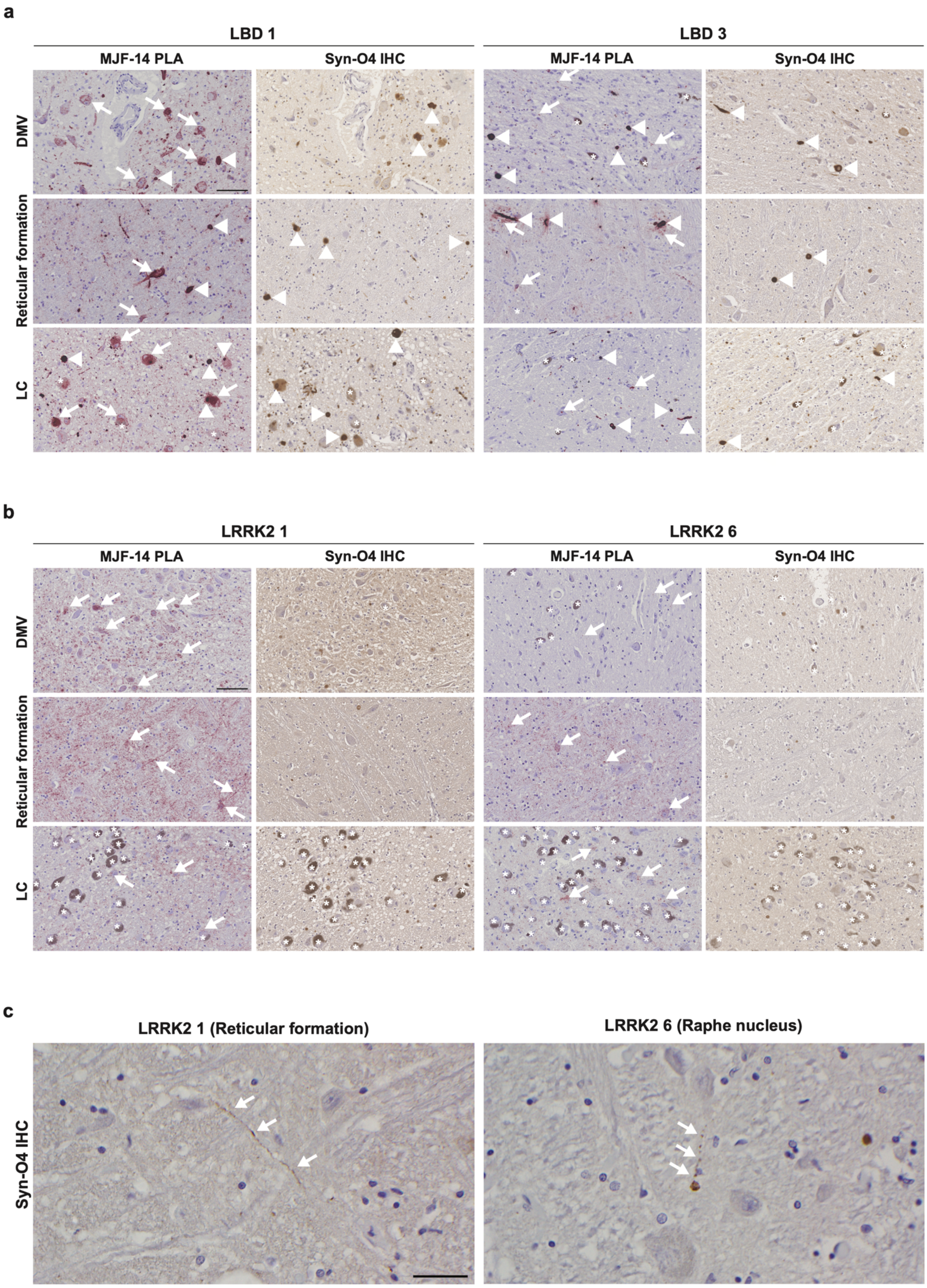
a
LBD 1
LBD 3
MJF-14 PLA
Syn-O4 IHC
MJF-14 PLA
Syn-O4 IHC
DMV
Reticular formation
LC
b
LRRK2 1
LRRK2 6
MJF-14 PLA
Syn-O4 IHC
MJF-14 PLA
Syn-O4 IHC
DMV
Reticular formation
LC
c
LRRK2 1 (Reticular formation)
LRRK2 6 (Raphe nucleus)
Syn-O4 IHC

**Suppl. Fig. 2 Comparison of MJF-14 PLA and Syn-O4 IHC in medullar and pontine regions**

**a**) Where Syn-O4 IHC primarily labels Lewy-type inclusions (LBs and LNs, arrowheads) in LBD cases, MJF-14 PLA reveals widespread accumulation of small non-inclusion aggregates. These aggregates are found in neuronal somas (arrows), but also widely distributed outside the cell bodies (presumably mostly in presynaptic and axonal compartments). **b**) In the LRRK2 cases, Syn-O4 IHC does not label distinctive $\alpha$-synuclein pathology, despite the abundance of particulate PLA signal (examples indicated by arrows). Scale bars = 100 µm (applies to all images). Neuromelanized neurons are indicated by asterisks. **c**) Rare Syn-O4-positive axonal varicosities (indicated by arrows) are found in some of the brainstem nuclei of LRRK2 1, LRRK2 2, and LRRK2 6. Scale bar = 50 µm (applies to both images). DMV = dorsal motor nucleus of the vagus, LC = locus coeruleus.

**Suppl. Fig. 3**

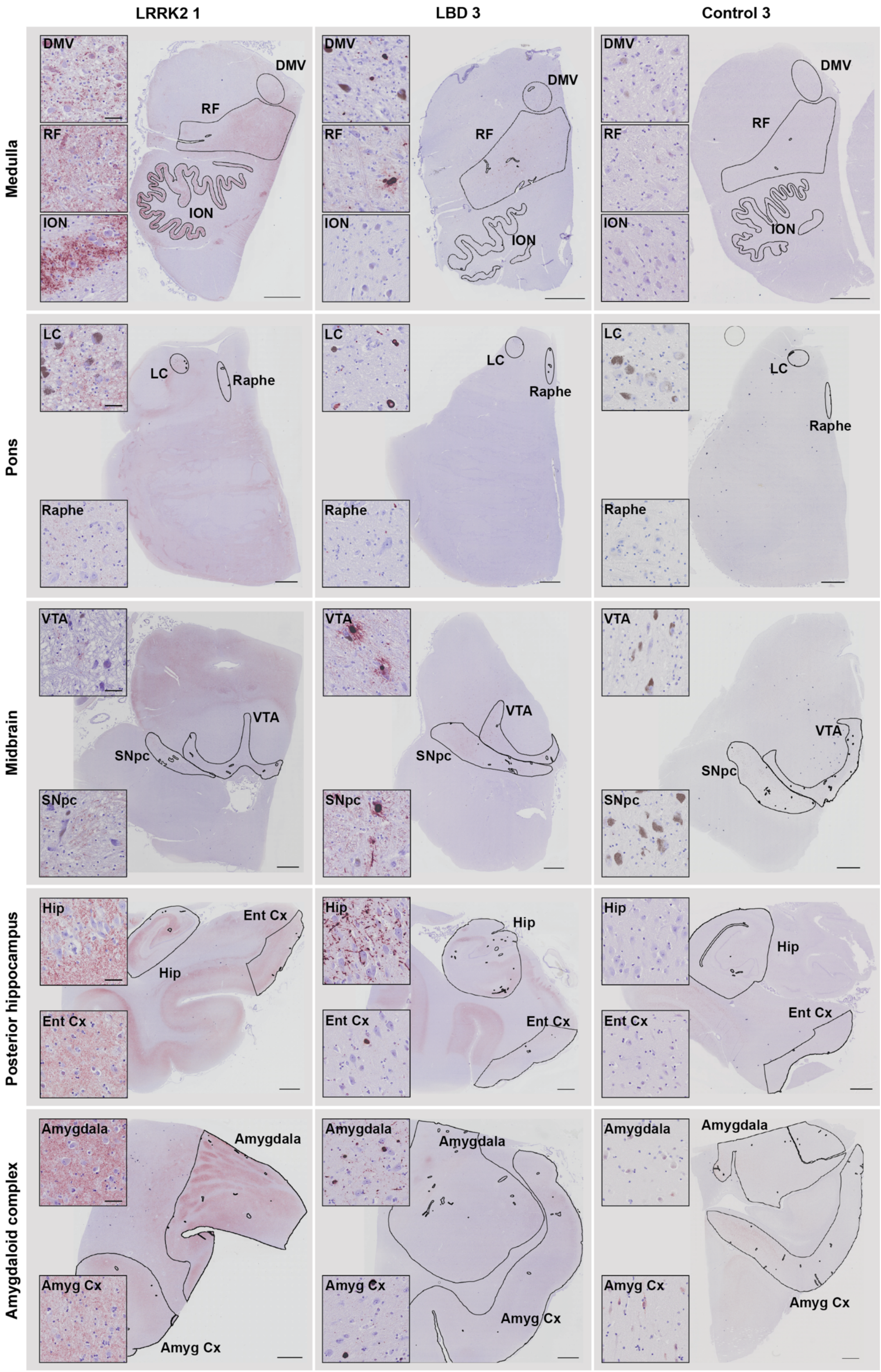

LRRK2 1
LBD 3
Control 3
Medulla
Pons
Midbrain
Posterior hippocampus
Amygdaloid complex
DMV
RF
ION
LC
Raphe
VTA
SNpc
Hip
Ent Cx
Amygdala
Amyg Cx

**Suppl. Fig. 3 Delineation of regions of interest for quantitative PLA analyses**

Full slide scans and magnified panels from three different cases and all five regions (medulla, pons, midbrain, posterior hippocampus, and amygdaloid complex) stained for each case. Regions of interest for quantitative analysis are outlined in black in each tissue section, while magnified panels show representative staining in the indicated regions. DMV = dorsal motor nucleus of the vagus, RF = reticular formation, ION = inferior olivary nucleus, LC = locus coeruleus, Raphe = raphe nucleus, SNpc = substantia nigra pars compacta, VTA = ventral tegmental area, Hip = hippocampus, Ent Cx = entorhinal cortex, Amyg Cx = Amygdaloid cortex. Scale bars = 2 mm (overview images), 50 µm (insets, applies to all images).

### Suppl. Fig. 4

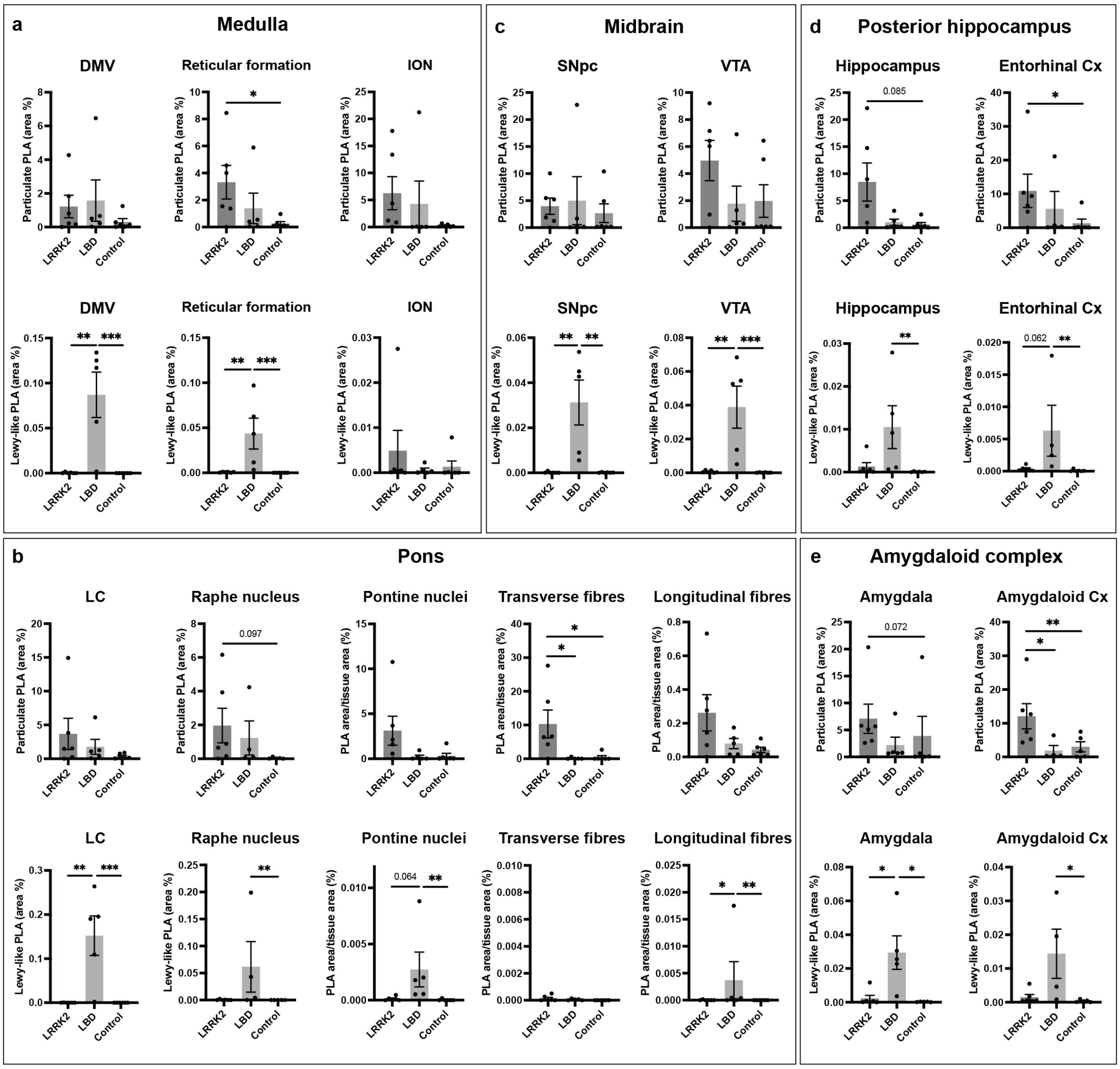

**Suppl. Fig. 4 Particulate PLA and Lewy-like PLA area coverage in various subregions**

Particulate PLA coverage (top) and Lewy-like PLA coverage (bottom) in subregions of the medulla, pons, midbrain, posterior hippocampus, and amygdaloid complex. All graphs show mean ± SEM with data points for individual cases. Univariate analyses covarying for age and sex followed by Bonferroni's multiple comparison correction. * p<0.05, ** p<0.01, *** p<0.001.

**Suppl. Fig. 5**

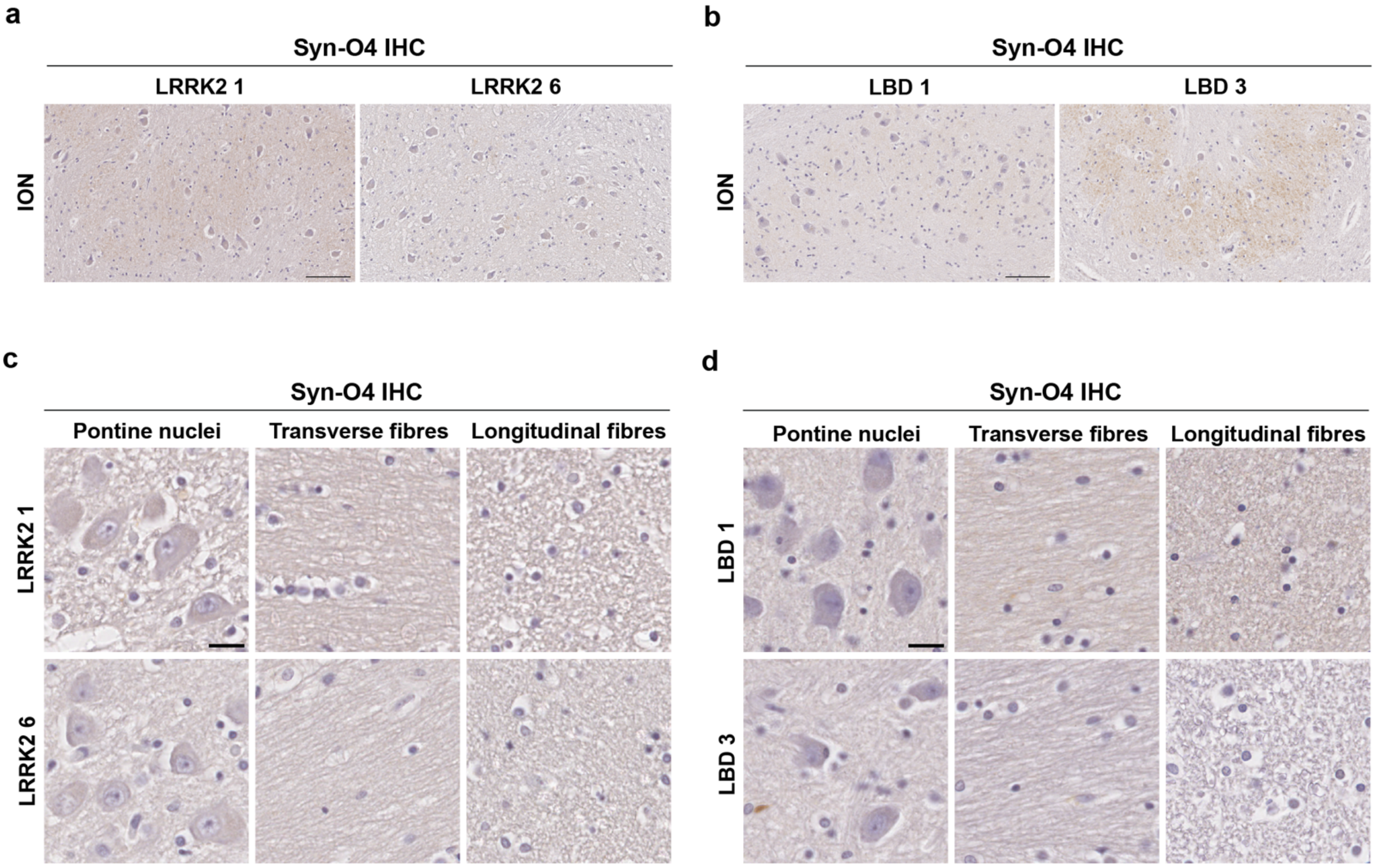

**Suppl. Fig. 5 Syn-O4 IHC in inferior olivary nucleus and basilar pons**

**a-b**) Syn-O4 IHC in the inferior olivary nucleus (ION) of two LRRK2 cases (**a**) and two LBD cases (**b**). No distinct signal corresponding to the particulate PLA signal in this region (see Fig. 4) is seen, though a weak brown background is seen in some cases. Note that the presence of this background does not appear to correspond to presence of PLA signal (more intense in LBD 3 than LBD 1, though LBD 1 has far more particulate PLA in ION). Scale bars = 100 µm (applies to all images). **c-d**) Syn-O4 IHC in pontine nuclei, transverse fibres, and longitudinal fibres of two LRRK2 cases (**c**) and two LBD cases (**d**). No distinct Syn-O4 signal is seen in these regions in any of the cases. Scale bars = 20 µm (applies to all images).

**Suppl. Table 1 Fold difference in area coverage of signal between particulate PLA and Lewy-like PLA for LBD cases**

| | LBD 1 | LBD 2 | LBD 3 | LBD 4 | LBD 5 | Average |
|---|---|---|---|---|---|---|
| **Medulla** | 121.09 | 10.85 | 12.40 | 56.59 | 7.26 | 41.64 |
| **Pons** | 39.76 | 51.87 | 14.09 | 184.51 | 12.89 | 60.62 |
| **Midbrain** | 272.23 | 34.23 | 24.36 | 18.87 | 5.64 | 71.07 |
| **Posterior hippocampus** | 983.56 | 242.41 | 286.16 | 115.90 | 41.10 | 333.83 |
| **Amygdaloid complex** | 174.80 | 239.65 | 28.66 | 46.65 | 14.84 | 100.92 |

**Suppl. Table 2 Reported neuropathological findings in LRRK2 mutation carriers**

| **Study**<br>Centre | **No. cases** | **Mutation and clinical diagnosis**[a] | **αSyn antibody** | **Pathological findings** | |
|---|---|---|---|---|---|
| | | | | **LB pathology** | **Non-LB pathology** |
| **Henderson et al. 2019** [32]<br>Center for Neuro-degenerative Disease Research, UPenn, US | 11 | 9 with G2019S mutation (7 with PD, 2 with PDD)<br>1 with L1165P (PDD)<br>1 with R793M (PDD) | pSer129 | **64% (7/11)**<br>prominent αSyn pathology (substantia nigra, amygdala, hippocampus and cingulate cortex but no staging given) | **36% (4/11)**<br>variable AD-like tau pathology |
| **Blauwendraat et al. 2019** [9]<br>Johns Hopkins Brain Resource Center, US | 6 | G2019S mutation (2 with PD, 3 with PDD, 1 with parkinsonism) | N/A | **33% (2/6)**<br>1 with neocortical LBD (+AD pathology)<br>1 with limbic LBD (+AD pathology) | **67% (4/6)**<br>3 with tau pathology (PSP-like)<br>1 with unspecific nigral cell loss |
| **Sanchez-Contreras et al. 2017** [68]<br>Mayo Clinic Florida brain bank, US | 4 | 1 with G2019S (probable PSP)<br>1 with R1441C (probable PSP)<br>1 with A1413T (PSP)<br>1 with R1707K (AD-type dementia) | N/A | -<br>(screened cohort only consisted of PSP and CBD) | **100% (4/4)**<br>2 pure PSP<br>1 PSP (+AD pathology)<br>1 CBD (+AD pathology) |
| **Ling et al. 2013** [44]<br>Queen Square Brain Bank, UCL, UK | 1 | G2019S mutation<br>and Q124E in MAPT (PD) | pSer129, KM51 | - | **100% (1/1)**<br>AD-type tau pathology (Braak tau stage III, CERAD B, Thal phase 3) and sparse TDP-43 pathology |
| **Poulopoulos et al. 2012** [59]<br>Columbia University Brain bank, New York, US | 3 | G2019S mutation (1 PD, 2 PDD) | N/A | **100% (3/3)**<br>1 with neocortical LBD (+AD pathology)<br>1 with neocortical LBD<br>1 brainstem LBD | - |
| **Ujiie et al. 2012**[b] [82]<br>Department of Pathology, Kanagawa, Japan | 6[b] | I2020T mutation (all with PD) | pSer129 | **17% (1/6)**<br>1 with LB pathology (distribution not reported) | **83% (5/6)**<br>2 with mild AD-type pathology (Braak stage 2 and 3)<br>2 with brainstem NFTs alone<br>1 with unspecific nigral loss |
| **Puschmann et al. 2012** [60]<br>Lund University, Sweden | 1 | N1437H mutation (PD) | LB509 | **100% (1/1)**<br>neocortical LBD with prominent ubiquitin pathology, also in white matter | - |
| **Ruffmann et al. 2012** [67]<br>Division of Pathology, Milan, Italy | 1 | G2019S mutation (PDD) | 4D6 | - | **100% (1/1)**<br>1 with tau pathology (PSP-like) |

| | | | | | |
|---|---|---|---|---|---|
| **Gomez and Ferrer 2010** [27]<br>Hospital Bellvitge, University of Barcelona, Spain | 3 | G2019S mutation (PD) | pSer129,<br>Chemicon,<br>Syn 505 | **67% (2/3)**<br>2 limbic DLB | **33% (1/3)**<br>Unspecific nigral cell loss |
| **Martí-Massó et al. 2009** [48]<br>Pathologic Anatomy Department, San Sebastian, Spain | 1 | R1441G mutation (PD) | pSer129, Novocastra,<br>Chemicon,<br>Syn 505 | - | **100% (1/1)**<br>Unspecific nigral cell loss |
| **Covy et al. 2009** [15]<br>Department of Pathology, UPenn, US | 2 | 1 with R793M mutation (PD)<br>1 with L1165P mutation (PDD) | pSer129,<br>LB509,<br>Syn 514,<br>Syn 21,<br>In-house SNL4 against aa 2-12 | **100% (2/2)**<br>Neocortical DLB with occasional TDP-43 pathology in temporal cortex | - |
| **Hasegawa et al. 2009**[b] [31]<br>Sagamihara National Hospital, Japan | 8[b] | I2020T mutation (PD) | pSer129 | **12% (1/8)**<br>1 with brainstem LBD | **88% (7/8)**<br>6 with unspecific nigral cell loss<br>1 with MSA-P type pathology |
| **Giordana et al. 2007** [26]<br>Department of Neuroscience, University of Torino, Italy | 1 | I1371V mutation (PD) | KM51,<br>LB509,<br>Syn1 | **100% (1/1)**<br>Limbic LBD (+AD pathology) | - |
| **Gaig et al. 2007** [22]<br>Hospital Clinic, University of Barcelona, Spain | 1 | G2019S mutation (PD) | N/A | - | **100% (1/1)**<br>Unspecific nigral cell loss |
| **Dächsel et al. 2007** [16]<br>Mayo Clinic Florida brain bank, US | 1 | G2019S mutation (no detailed clinical data but dementia and tremor) | N/A | - | **100% (1/1)**<br>Ubiquitin-IR inclusion alone, similar to FTLD-U |
| **Giasson et al. 2006** [24]<br>Department of Pathology, UPenn, US | 3 | G2019S mutation (parkinsonism) | LB509,<br>Syn 505 | **67% (2/3)**<br>1 with neocortical LBD (+AD pathology)<br>1 with brainstem LBD | **33% (1/3)**<br>1 with unspecific nigral cell loss |
| **Ross et al. 2006** [65]<br>Mayo Clinic Florida brain bank, US | 8 | G2019S mutation (PD) | N/A | **100% (8/8)**<br>4 with brainstem LBD<br>3 with limbic LBD<br>1 with neocortical LBD | - |
| **Rajput et al. 2006** [61]<br>Royal University Hospital, Saskatoon, Canada | 3 | G2019S mutation (PD) | LB509 | - | **100% (3/3)**<br>2 with tau pathology (PSP-like)<br>1 with argyrophilic grain disease |

| | | | | | |
|---|---|---|---|---|---|
| **Khan et al. 2005** [40]<br>Queen square Brain Bank, UCL, UK | 1 | Y1699C mutation (PD) | N/A | **100% (1/1)**<br>Brainstem LBD | - |
| **Gilks et al. 2005** [25]<br>Queen square Brain Bank, UCL, UK | 3 | G2019S mutation (PD) | N/A | **100% (3/3)**<br>1 with brainstem LBD<br>2 with limbic LBD | - |
| **Wszolek et al. 2004**[c] [87]<br>Mayo Clinic Florida brain bank, US | 4[c] | R14441C mutation (3 with PD and 1 with PSP) | N/A | **50% (2/4)**<br>1 with brainstem LBD<br>1 with neocortical LBD | **50% (2/4)**<br>1 with PSP-like tau pathology<br>1 with unspecific nigral cell loss |
| **Zimrich et al. 2004**[c] [90]<br>Mayo Clinic Florida brain bank, US | 4[c] | R14441C mutation (3 with PD and 1 with PSP) | N/A | **50% (2/4)**<br>1 with brainstem LBD<br>1 with neocortical LBD | **50% (2/4)**<br>1 with PSP-like tau pathology<br>1 with unspecific nigral cell loss |
| **Wszolek et al. 1997** [88]<br>University of Nebraska Medical Center, Omaha, US | 2 | Y1699C mutation (PD) | None (Ubiquitin) | - | **50% (1/2)**<br>Both cases had nigral cell loss without LB pathology (one with AD pathology) |
| **TOTAL** | **68** | | | **33 (48.5%)** | **35 (51.5%)** |

Abbreviations: AD = Alzheimer's disease, CBD = corticobasal degeneration, FTLD-U = frontotemporal lobar degeneration with ubiquitin-positive inclusions, LBD = Lewy body disease, MSA-P = multiple system atrophy parkinsonism-subtype, NFTs = neurofibrillary tangles, PSP = progressive supranuclear palsy.

[a] Unless otherwise stated, mutations listed all pertain to LRRK2. Where available, clinical diagnosis is listed in brackets.

[b] These two studies have same cases.

[c] These two studies have same cases.
